# Comparative Multiomic Analysis Reveals Low T Cell Infiltration as the Primary Feature of Tobacco Use in HPV(+) Oropharyngeal Cancer

**DOI:** 10.1101/2021.03.23.436478

**Authors:** Benjamin M. Wahle, Paul Zolkind, Ricardo Ramirez, Zachary L. Skidmore, Angela Mazul, D. Neil Hayes, Vlad C. Sandulache, Wade L. Thorstad, Douglas Adkins, Obi L. Griffith, Malachi Griffith, Jose P. Zevallos

**Affiliations:** Washington University Department of Otolaryngology – Head and Neck Surgery, St. Louis, Missouri, USA; McDonnell Genome Institute, Washington University School of Medicine, St. Louis, Missouri, USA; Department of Medicine, Division of Hematology-Oncology, University of Tennessee Health Science Center, Memphis, Tennessee, USA; Bobby R. Alford Department of Otolaryngology Head and Neck Surgery, Baylor College of Medicine, Houston, Texas, USA; ENT Section, Operative Care Line, Michael E. DeBakey Veterans Affairs Medical Center, Houston, Texas, USA; Department of Radiation Oncology, Washington University School of Medicine, St Louis, MO, United States; Department of Medicine, Division of Oncology, Washington University School of Medicine, St. Louis, Missouri, USA

## Abstract

**Purpose:** Tobacco use is an independent adverse prognostic feature in human papillomavirus (HPV)-associated oropharyngeal squamous cell carcinoma (OPSCC). Despite this, the biologic features associated with tobacco use have not been systematically investigated in this population. We sought to characterize the genomic and immunologic features of HPV(+) OPSCC associated with tobacco use and adverse oncologic outcomes.

**Experimental Design:** Whole exome sequencing of 47 primary HPV(+) OPSCC tumors was performed to investigate mutational differences associated with tobacco exposure. To characterize the tumor immune microenvironment (TIME), targeted mRNA hybridization was performed and immunohistochemical (IHC) staining was used to validate these findings.

**Results:** Low expression of transcripts in a T cell-inflamed gene expression profile (TGEP) was associated with tobacco use at the time of diagnosis and lower overall and disease-free survival. Tobacco use was associated with an increased proportion of T>C substitutions and a lower proportion of mutational signatures typically observed in HPV(+) OPSCC tumors, but was not associated with increases in mutational burden or the rate of recurrent oncogenic mutations.

**Conclusions:** In HPV(+) OPSCC, low T cell infiltration of primary tumors is associated with current tobacco use and worse oncologic outcomes. Rather than an increased mutational burden, tobacco’s primary and clinically relevant association is immunosuppression of the primary TIME. An objective clinical assay like the TGEP, which quantifies immune infiltration of the primary TIME, may have value for HPV(+) OPSCC risk stratification in future clinical trials.

## INTRODUCTION

Oropharyngeal squamous cell carcinoma (OPSCC) is among the most common cancers of the upper aerodigestive tract. An estimated 60-70% of new OPSCC diagnoses are attributable to oncogenic human papillomavirus (HPV) exposure.^1^ The incidence of HPV(+) OPSCC is rising, and OPSCC has surpassed cervical cancer as the most common HPV-related malignancy in the United States.^1–3^

HPV(+) OPSCC is both molecularly and clinically distinct from HPV(−) disease, for which tobacco and/or alcohol use are the key causative exposures. While HPV(−) OPSCC often portends a grim prognosis, HPV(+) OPSCC outcomes are comparably favorable, with approximately 75-80% of HPV(+) OPSCC patients surviving five years after diagnosis.^2,4^ However, for most patients who respond to treatment, the morbidity associated with surgery, radiation therapy (RT), and/or platinum-based chemotherapy is substantial and often lifelong. The extent to which this treatment morbidity is justified by favorable oncologic outcomes is not clear. Treatment deintensification for this patient group is a major focus of recent and ongoing clinical trials in HPV(+) OPSCC.^5^ The unique challenges in the future of HPV(+) OPSCC treatment are safely reducing iatrogenic morbidity through treatment deintensification in low-risk individuals while also improving outcomes in the high-risk patients with aggressive disease not cured with current standard of care therapies. Importantly, both tasks rely on accurately predicting high or low-risk disease.

Tobacco use is well-established as an independent adverse prognostic factor in HPV(+) OPSCC.^2,6–10^ Multiple reports have demonstrated that tobacco use is associated with increased risk of disease-specific adverse outcomes, suggesting that tobacco users’ attenuated prognosis is not entirely attributable to competing causes of mortality.^6,7,10,11^ Tobacco users with HPV(+) OPSCC represent an intermediate prognostic group, between tobacco-naïve HPV(+) patients with an excellent prognosis and HPV(-) OPSCC patients whose prognosis is poor.^2,6,10^ Despite this, the molecular features associated with tobacco use in HPV(+) OPSCC are poorly defined. We hypothesized that tobacco influences two biologic processes which could contribute to more aggressive disease and worse outcomes in HPV(+) OPSCC. First, tobacco may be associated with increased somatic mutational burden and/or a distinct array of oncogenic mutations, contributing to more aggressive disease. Despite tobacco’s canonical role as a mutagen in head and neck squamous cell carcinoma (HNSCC), it has not been clearly demonstrated that tobacco exposure influences mutational burden when HPV is the causative exposure.^12^ Second, tobacco exposure may be associated with changes to the tumor immune microenvironment (TIME) that produce relative immunosuppression and a more aggressive disease course. This hypothesis is supported by a recent report demonstrating lower CD8+ immunohistochemical (IHC) staining in tobacco-exposed HPV(+) tumors.^13^ No prior investigation has systematically evaluated the molecular features associated with tobacco, comparing both its mutational and tumor immune effects. Using a multi-omics approach, we tested both of these hypotheses in a cohort of HPV(+) OPSCC patients. We evaluated somatic mutational burden with whole exome sequencing (WES) and investigated the TIME by measuring expression of immune-related mRNA transcripts and IHC staining.

## METHODS

### Cohort and sample acquisition

All patients were treated at the Washington University Siteman Cancer Center. After informed consent, samples were collected prospectively from patients with newly diagnosed, biopsy proven, p16-positive OPSCC as part of the Washington University tumor banking protocol. The tumor acquisition protocol and correlative studies were approved by the Washington University Human Research Protection Office. Demographic and clinical data were collected from medical records. Tobacco use information was available for all patients, who were assigned to current, former, or never tobacco use categories. Dates of diagnosis, death, or recurrence (local, regional, or distant) were recorded for each patient and used to calculate overall and disease-free survival (OS and DFS, respectively).

### Sequencing

Genomic DNA was isolated from 47 paired tumor/normal samples using Qiagen DNeasy Blood and Tissue Kits. Genomic DNA was fragmented to 100-300bp and libraries were constructed using the KAPA HTP Library Kits (KAPA Biosystems). A single low input sample used the Swift Accel-NGS 2S DNA Library Kit (Swift BioSciences) for library construction. Samples were then sequenced to a target depth of 120x per sample on the Illumina NovaSeq platform (S4, 2 × 151 bp reads). Additional computational methods are detailed in the **Supplementary Methods**.

### NanoString mRNA hybridization

RNA was isolated from 47 tumor samples using QIAGEN RNeasy kits. Transcripts were hybridized to a panel of 770 probes (NanoString IO360). Hybridized transcripts were purified and immobilized on a sample cartridge. Counts of hybridized transcripts were read using the nCounter platform. Raw counts were normalized with the nSolver software (4.0) package using the geometric mean of both positive control probes and housekeeping probes. A background threshold count was set to 20. Normalized data was used for hierarchical clustering. Over representation analysis (ORA) was performed using WebGestalt.^14^ T cell-inflamed gene expression profile (TGEP) scores were calculated separately by normalizing using all available housekeeping probes and weighting transcript values as previously described.^15,16^ Validation of TGEP score findings was performed by identifying p16+ OPSCC cases from the HNSC-TCGA with available normalized RNAseq data. For a cross-platform comparison, z-scores were calculated for each of the genes in the TGEP score and unsupervised clustering was performed, followed by annotation with clinical data.

### IHC

Tumor microarrays were assembled from formalin-fixed paraffin-embedded primary tumors. A clinical pathologist reviewed slides and selected areas containing tumor for inclusion in cores. Slides of 5 micron thickness were made and stained with Ventana antibodies for CD3 (2GV6), CD4 (SP35), CD8(SP57), and PD-L1(SP263) with hematoxylin counterstaining. A high-resolution digital scanner (NanoZoomer; Hamamatsu Photonics, Welwyn Garden City, UK) was used to create digitized bright field images from the slides at 20x magnification which spanned the entire area of each core. Cores were not included in analyses if >30% of the core was missing or damaged during sectioning. The images were evaluated using digital analysis software (Visiopharm; Hoersholm, Denmark), which has been used previously for identifying T cells in the HNSCC TIME.^17^ Red-Blue-Green color filters were set which reliably distinguished 3,3’-diaminobenzidine (DAB) stained and hematoxylin stained cells without detecting unstained negative space. Areas of artifact or damage were manually excluded. The total area stained by DAB or hematoxylin was calculated per core. Positive staining for each core was quantified and reported as the total DAB[+] area divided by the total cellular area (DAB[+] plus hematoxylin[+]). Example images are shown for anti-CD8 stained tumors in **Figure S7**.

### Statistical Methods

Statistical analyses were performed in R (3.6.1) or GraphPad Prism (8.1.2). Shapiro Wilk tests were used to determine normality of distributions. Normally distributed continuous variables were compared using two-sided T-tests with Welch’s correction. Variables without normal distributions were compared with Mann Whitney or Kruskal Wallis tests with Dunn’s multiple comparison tests. Spearman’s rank correlation coefficients were computed to determine degrees of correlation. Contingency testing was performed using Fisher’s exact tests. Survival between groups was compared using Log-rank tests. False Discovery Rates were computed when multiple comparisons were performed together. Cut point analysis was performed using the R package survminer (0.4.2). Hierarchical clustering was performed using Euclidian distance with Ward’s linkage method.

## RESULTS

### Cohort characteristics

Demographic and clinical characteristics of our cohort are displayed in **Table 1**. The majority of patients had a history of tobacco exposure, with 9 (19.1%) being current users, 19 (40.4%) being former users, and 19 (40.4%) being never users. Consistent with established demographic trends in HPV(+) OPSCC, the median age at diagnosis was 58 years old (range 45 – 79) and the majority of patients were White males. No significant differences in demographic and American Joint Committee on Cancer (AJCC) 8^th^ edition staging parameters were present based on tobacco exposure groups. Our cohort included patients treated with primary surgery (76.6%) or definitive chemoradiation therapy (CRT) (23.4%). The median time to loss of follow up or death was 1.9 years (range 0.23 to 7.8 years). Eight patients (17.0%) in the cohort had disease recurrence and six (12.8%) died during the study period. OS and DFS were both significantly associated with clinical disease stage (**Supplemental Figure S1A,B**; both P < 0.001). There were no OS or DFS differences related to tobacco use status (**Supplemental Figure S1C,D**; P = 0.220 and 0.650, respectively).

**Table 1:** Demographic and clinical characteristics of the study cohort. All patients were p16(+) and had untreated primary OPSCC.

|  | Current<br>(N=9) | Former<br>(N=19) | Never<br>(N=19) | P value | Total<br>(N=47) |
| --- | --- | --- | --- | --- | --- |
| **Age at Diagnosis** |  |  |  |  |  |
| Mean (SD) | 57.6 (5.22) | 61.3 (10.6) | 56.9 (9.07) | 0.312 | 58.8 (9.26) |
| **Decade of Diagnosis** |  |  |  |  |  |
| 50-59 | 6 (66.7%) | 7 (36.8%) | 7 (36.8%) | 0.212 | 20 (42.6%) |
| 60-69 | 3 (33.3%) | 5 (26.3%) | 4 (21.1%) |  | 12 (25.5%) |
| 40-49 | 0 (0%) | 2 (10.5%) | 6 (31.6%) |  | 8 (17.0%) |
| 70-79 | 0 (0%) | 5 (26.3%) | 2 (10.5%) |  | 7 (14.9%) |
| **Gender** |  |  |  |  |  |
| Female | 1 (11.1%) | 3 (15.8%) | 1 (5.3%) | 0.828 | 5 (10.6%) |
| Male | 8 (88.9%) | 16 (84.2%) | 18 (94.7%) |  | 42 (89.4%) |
| **Race/Ethnicity** |  |  |  |  |  |
| Asian | 0 (0%) | 1 (5.3%) | 0 (0%) | 1 | 1 (2.1%) |
| Caucasian | 9 (100%) | 18 (94.7%) | 19 (100%) |  | 46 (97.9%) |
| **Primary Tumor Subsite** |  |  |  |  |  |
| Base of Tongue | 3 (33.3%) | 9 (47.4%) | 8 (42.1%) | 0.838 | 20 (42.6%) |
| Tonsil | 6 (66.7%) | 9 (47.4%) | 11 (57.9%) |  | 26 (55.3%) |
| Overlapping Sites | 0 (0%) | 1 (5.3%) | 0 (0%) |  | 1 (2.1%) |
| **Clinical T Stage** |  |  |  |  |  |
| T0 | 1 (11.1%) | 2 (10.5%) | 0 (0%) | 0.462 | 3 (6.4%) |
| T1 | 2 (22.2%) | 4 (21.1%) | 3 (15.8%) |  | 9 (19.1%) |
| T2 | 2 (22.2%) | 7 (36.8%) | 12 (63.2%) |  | 21 (44.7%) |
| T3 | 3 (33.3%) | 3 (15.8%) | 3 (15.8%) |  | 9 (19.1%) |
| T4 | 1 (11.1%) | 3 (15.8%) | 1 (5.3%) |  | 5 (10.6%) |
| **Clinical N Stage** |  |  |  |  |  |
| N0 | 1 (11.1%) | 2 (10.5%) | 0 (0%) | 0.166 | 3 (6.4%) |
| N1 | 4 (44.4%) | 5 (26.3%) | 10 (52.6%) |  | 19 (40.4%) |
| N2 | 3 (33.3%) | 12 (63.2%) | 7 (36.8%) |  | 22 (46.8%) |
| N3 | 1 (11.1%) | 0 (0%) | 2 (10.5%) |  | 3 (6.4%) |
| **Clinical M Stage** |  |  |  |  |  |
| M0 | 9 (100%) | 18 (94.7%) | 19 (100%) | 1 | 46 (97.9%) |
| M1 | 0 (0%) | 1 (5.3%) | 0 (0%) |  | 1 (2.1%) |
| **Overall Clinical Stage** |  |  |  |  |  |
| I | 3 (33.3%) | 4 (21.1%) | 8 (42.1%) | 0.658 | 15 (31.9%) |
| II | 4 (44.4%) | 12 (63.2%) | 8 (42.1%) |  | 24 (51.1%) |
| III | 2 (22.2%) | 2 (10.5%) | 3 (15.8%) |  | 7 (14.9%) |
| IV | 0 (0%) | 1 (5.3%) | 0 (0%) |  | 1 (2.1%) |

WES of tumor-normal pairs was performed for all 47 patients in our cohort, with a median depth of sequencing of 71x for tumor and 95x for normal samples. Prior to testing our hypotheses related to tobacco exposure, we determined the recurrent mutations and structural alterations within the cohort. We identified 12 genes that met criteria for statistically significant mutations in our cohort (MuSiC FDR < 0.1). These included multiple genes identified in HPV(+) OPSCC in previous publications,^12,18^ including *PIK3CA, ZNF750, FGFR3, PTEN, TRAF3,* AND *FBXW7* (**Figure 1; Supplemental Tables S1A,B**). Additionally, this approach identified mutations in genes that have not previously been noted as statistically significant in HPV(+) OPSCC, including *B2M, IFI27, AK5, METTL24, IQCG,* and *SMARCAL1.* Using a classifier which considers mutations of a gene as well as mutations in the particular gene’s functionally related network neighbors,^19^ we identified additional recurrent mutations in *AKT1, CUX1, FGF2, FGF8, HLA-B, KMT2D, NRAS, PIK3R1, PLEC, SYNE1,* and *SYNE2* (MUFFINN probabilistic score >0.5, **Supplemental Table S1C**).

**Figure 1:**
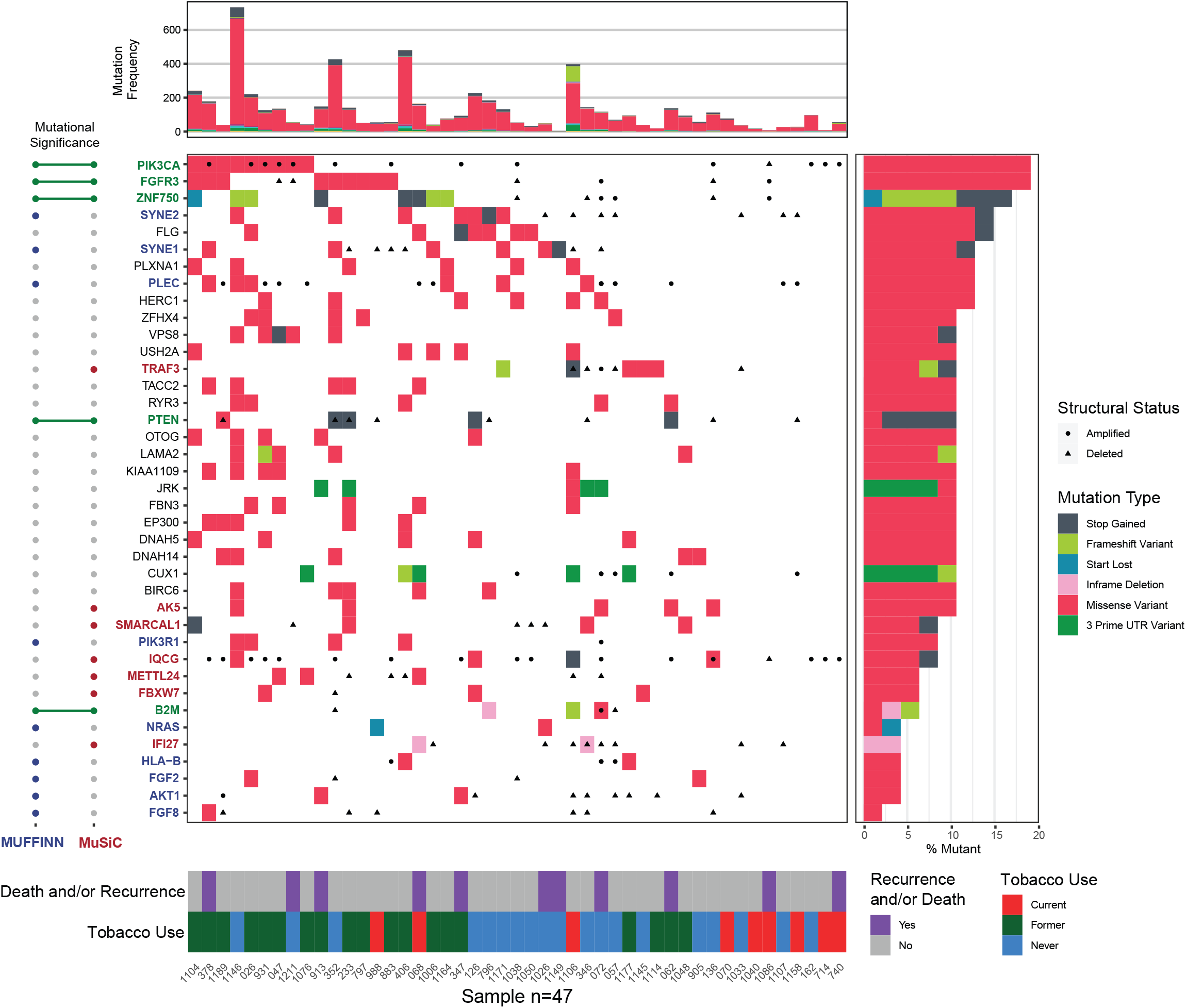
Somatic mutations in HPV(+) OPSCC. *Top panel:* Mutational frequency is displayed for each patient, representing the total number of nonsynonymous single nucleotide variations after filtration. Left panel: Mutated genes are ranked from most frequently mutated (top) to least (bottom) and are annotated by mutational significance classifier. Genes in our cohort not previously noted as statistically significantly mutated include *B2M, SMARCAL1, AK5, IQCG, METTL24, and IFI27*. *Center panel:* Waterfall plot of somatic mutations and mutational consequences for each patient. Focal copy number changes at each locus were included for those with a log2 ratio greater than +/− 0.5. *Right panel:* Frequency and classifications of somatic mutations for each gene. *Bottom panel:* Clinical annotation

Many observed copy number alterations (CNAs) were consistent with previous reports and included amplifications on 1q, 3q, 5p, 8q, 12p, 19q, and 20q and deletions on 3p, 9q, 11q, 14q, and 16q (**Supplemental Figure S2**).^12,18^ Multiple focal amplifications and deletions not described for HPV(+) OPSCC in previous publications met criteria for statistical significance (q < 0.01). The most notable were an amplification of 7p22.1 containing *RAC1* and a deletion of 13.q14.2 containing *RB1* (**Supplemental Table S1D,E**).

### Tobacco use is associated with differences in single base substitution signatures, but not mutational burden or recurrent oncogenic mutations

Because of tobacco’s causative role in HNSCC at multiple sites, we hypothesized that WES would reveal increased somatic mutational burden in patients with tobacco use history. Despite wide variability in the tumor mutational burden (TMB) throughout the cohort, ranging from 0.04 to 30.1 mutations/Mb, there were no significant differences in TMB related to history of tobacco use regardless of how we categorized tobacco use. We observed no difference in TMB comparing tobacco exposed versus never exposed users (P = 0.614), current versus former plus never tobacco users (P = 0.642) or by comparing current, former, and never tobacco use groups separately (**Supplemental Figure S3A**, P = 0.655). Additionally, TMB did not significantly differ in patients who had adverse clinical outcomes including death and/or recurrence (**Supplemental Figure S3B**, P = 0.738).

We sought to identify differences in mutational profiles based on tobacco exposure. In a subset of patients with sufficiently high mutational burden (total nonsynonymous SNVs >45, N = 39 patients, 83.0% of cohort), we examined the rate of single nucleotide substitutions and their trinucleotide contexts. Consistent with previous reports of HPV(+) OPSCC, tumors were dominated by C>T transitions and C>G transversions (**Figure 2A**).^12^ Tobacco-exposed tumors had a 2.3-fold greater median proportion of T>C transitions compared to tumors of never tobacco users (**Figure 2F**, P = 0.015, FDR = 0.088). Tobacco exposure was not associated with significant differences in the remaining substitution classes (**Figure 2B-E, G**).

**Figure 2:**
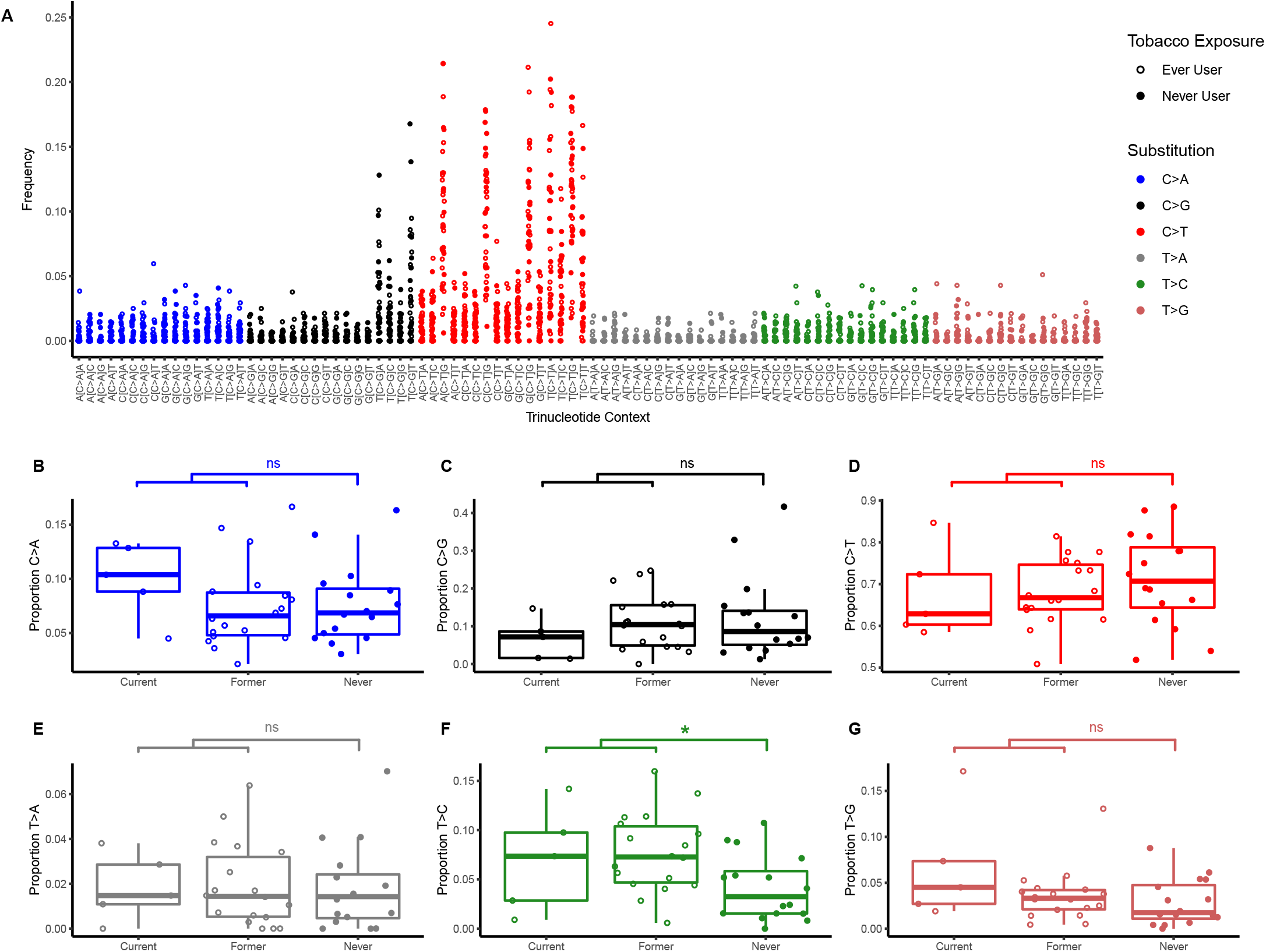
Tobacco exposure is associated with differences in trinucleotide substitution classes. **(A)** Consistent with previous reports, mutations in our cohort predominantly consisted C>T transitions and C>G transversions. **(B - G)** Each class of nucleotide substitution is displayed as a proportion of total mutations. **(F)** Tobacco exposed tumors demonstrated a 2.3-fold greater median proportion of T>C transitions compared to tumors of never tobacco users. (* P < 0.05 and FDR < 0.1, ns = not significant)

Single base substitution signatures were determined for each sample and compared based on tobacco use status.^20^ Consistent with prior reports, three mutational signatures were dominant (**Figure 3A**). Signature #1 was present in 92.3% of samples and reflects spontaneous 5-methylcytosine deamination associated with aging. Signatures #2 and #13, which reflect mutations attributable to APOBEC activity, were present in 74.3% of patients. The median proportion of mutations accounted for by signatures #1, #2, and #13 combined was 66.8% (range 9.2 to 91.7%). Together, these dominant signatures accounted for a significantly higher proportion of mutations in tumors of never tobacco-users when compared with tobacco exposed tumors (**Figure 3B**, P = 0.026).

**Figure 3:**
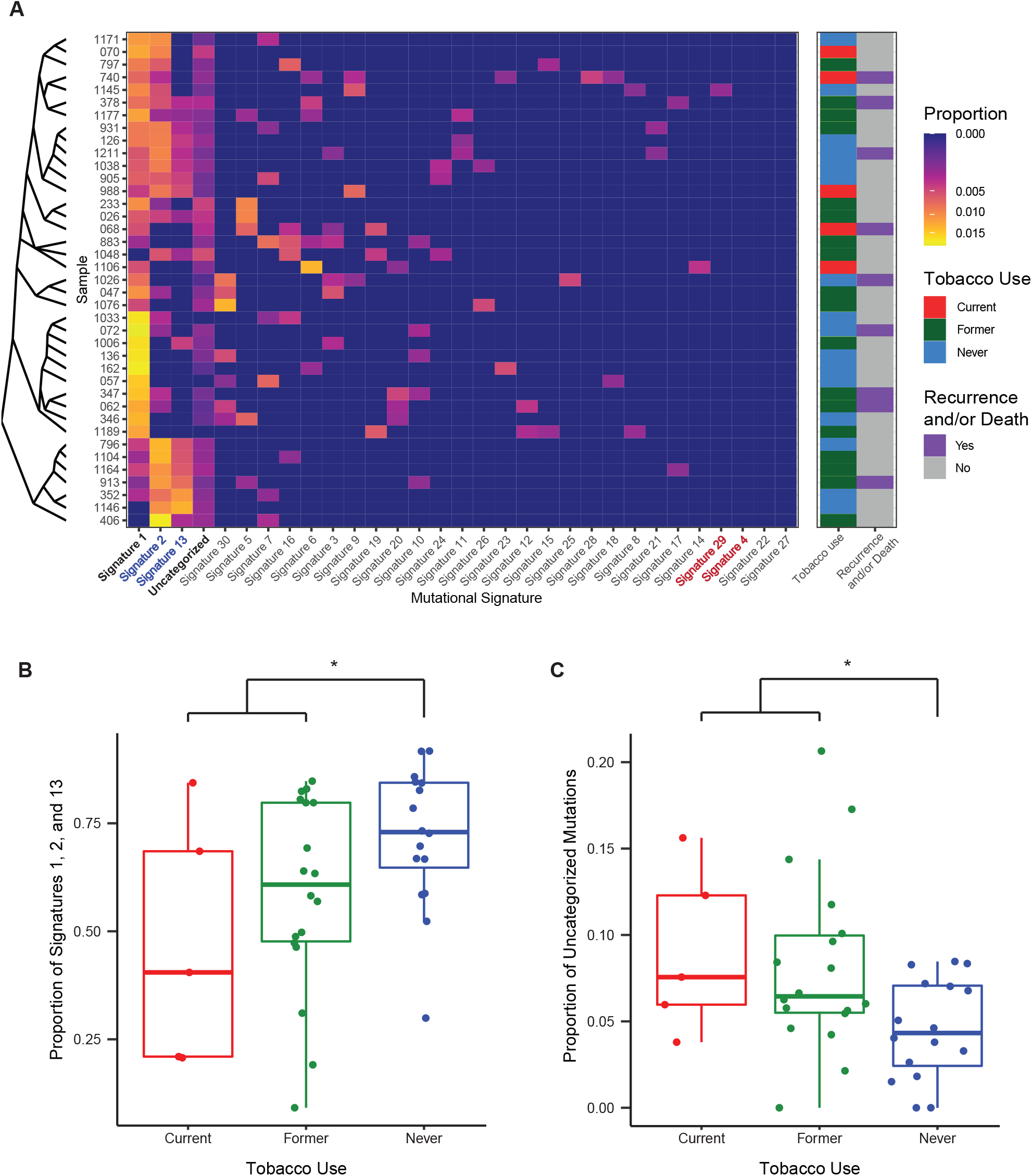
Aging and APOBEC-related mutational signatures are dominant in HPV(+) OPSCC but account for a lower proportion of mutations in tobacco users. **(A)** Heatmap of mutational signatures demonstrates the dominance of signatures #1 (Aging-related), #2 and 13 (both APOBEC-related) in HPV(+) OPSCC. Despite high reported rates of tobacco use in the cohort, mutations attributable to tobacco-related signatures #4 and #29 were absent. **(B)** The proportion of signature #1, #2, and #13 mutations is significantly greater in never tobacco users, while **(C)**the proportion of uncategorized mutations is significantly greater in those with a tobacco use history. (* P < 0.05)

Tobacco-related mutational signatures were largely absent within our cohort. Signature #4, which is associated with tobacco use, was not responsible for any proportion of mutations in our study population. Signature #29, associated with chewing tobacco use, was present in one patient accounting for 8.2% of mutations in this individual. Signature #16 mutations have been previously correlated with tobacco exposure in HPV(-) disease,^12^ but were not significantly associated with tobacco use (P = 0.418) in this cohort. We also considered the proportion of mutations that could not be categorized into one of the thirty mutational signatures. Uncategorized mutations were present in 94.9% of samples and accounted for a minority of mutations in most patients (median 6.0% of mutations, range 0 to 20.6%). Uncategorized mutations accounted for a significantly greater proportion of mutations in tobacco exposed tumors (**Figure 3C**, P = 0.017). This difference approached statistical significance with current, former, and never tobacco users compared separately (P = 0.052).

In an exploratory analysis, we compared the frequency of recurrent somatic mutations in the whole cohort by tobacco use groups. There were no significant differences in the occurrence of any single mutated gene based on tobacco use groups (**Table S1F**). We hypothesized that mutations typically observed in HPV(-) HNSCC may be induced by tobacco use. To test this, we examined the rate of canonical HPV(-) HNSCC mutations which were significantly mutated in TCGA(Supplemental **Table S1G**).^18^ These mutations individually occurred at a low rate in our cohort with no single gene having more than an 8.5% mutation rate. Collectively, canonical HPV(-) mutations occurred at a rate of 32.1% of those with a history of tobacco use compared to 15.8% of those with no tobacco history (P = 0.31).

### Targeted gene expression profiling reveals that T cell-inflamed TIME is inversely associated with current tobacco use

We hypothesized that tobacco use would be associated with discernible differences in the TIME. We utilized a 760-gene NanoString immuno-oncology panel to measure expression of immune-cell related transcripts. Of the 47 tumors available for analysis, three samples were excluded because they did not pass quality control review due to low binding density and/or low efficiency hybridization.

We performed unsupervised hierarchical clustering of the 760 transcripts for all patients. This approach divided patients into two clusters which differed by disease-free survival (**Figure 4**). A cluster of 10 low-risk patients was identified, none of which had death or recurrence, with all adverse events concentrated in the remaining 34 patients (P = 0.069, HR = 3.703, 95% CI 0.905-15.15). We observed that the 10 patient low-risk cluster was enriched for a cluster of 202 genes containing T cell related transcripts (**Figure 4**, column cluster 4), including T cell related cell surface markers, members of the T cell receptor complex, granzymes, and IFNλ and JAK-STAT pathway members (**Supplemental Table S2D**). ORA revealed that relative to all transcripts in the panel, this cluster was significantly enriched for a number of biologic processes related to adaptive immunity in all functional databases we queried (**Supplemental Table S2D**). IHC for CD3, CD4 and CD8 showed that the 10-patient low-risk cluster had a significantly greater degree of T cell infiltration (**Supplemental Figure S4A-C**, all FDR <0.01), confirming that enrichment for adaptive immune-related transcripts in this cluster correlates with protein expression of T cell surface markers.

**Figure 4:**
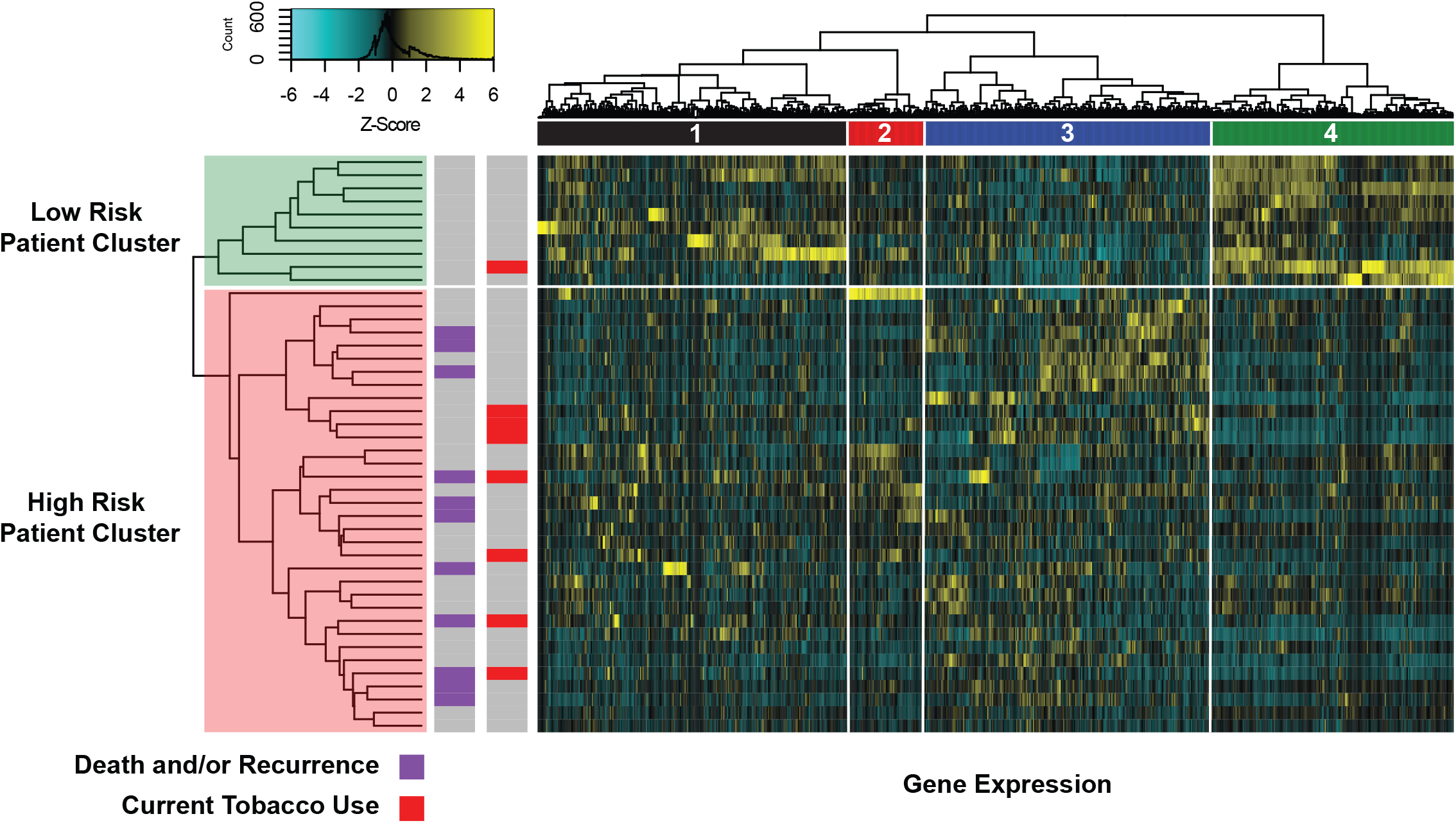
Unsupervised clustering of 760 transcripts reveals a low-risk patient group with high expression of T cell-related transcripts. Heatmap of gene expression displays high expression of transcripts in column cluster 4 in a subset of patients without death and/or recurrence. ORA of transcripts in this cluster revealed that cluster 4 was significantly enriched for multiple of biologic processes related to adaptive immunity. Patients in the high-risk patient cluster, who had lower expression of cluster 4 transcripts, displayed increased risk of death and/or recurrence (HR = 3.703, 95% CI 0.905-15.15, P = 0.069).

To further characterize the functional significance of T cell related differences suggested by unsupervised clustering, we utilized a previously published and validated 18-gene T cell-inflamed gene expression profile (TGEP) score developed on the NanoString platform.^15,16^ A high score reflects existing cytotoxic activity in the TIME and has been validated in multiple tumor types, including large cohorts of HNSCC patients, as a predictor of response to PD-1 inhibitors.^15^ **Figure 5A** displays the spectrum of transcript expression contributing to TGEP scores in our cohort. While TGEP scores showed significant positive correlations with IHC expression of CD3+, CD4+ and PD-L1 (**Supplemental Figure S5A,C,G**, all FDR < 0.1), the strongest and most significant correlation was with CD8+ cells (**Figure 5B**, Rho = 0.728, FDR < 0.001), suggesting that TGEP scores serve as a reliable indicator of a CD8+ T cell-inflamed TIME.

**Figure 5:**
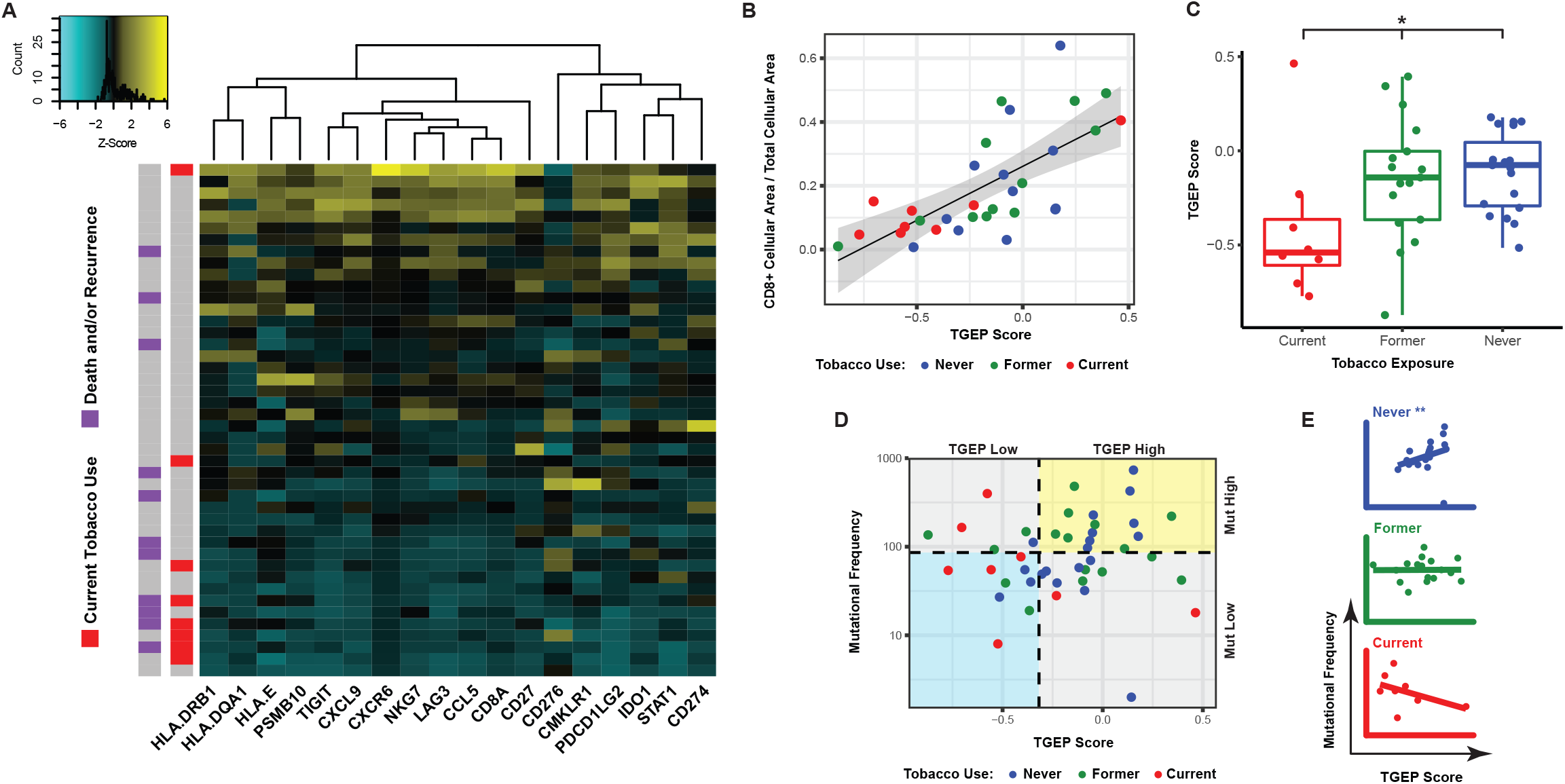
Tobacco use at the time of diagnosis is associated with decreased T cell infiltration of the primary TIME. **(A)** Heatmap displaying TGEP transcript expression for each patient. Patients are ranked from high (top) to low (bottom) based on their TGEP scores. **(B)** TGEP scores were significantly associated with CD8+ IHC staining (Rho = 0.728, P < 0.001). **(C)** TGEP scores differ significantly by tobacco use, with current tobacco users having the lowest scores. **(D)** Tobacco history reveals differences in TGEP scores (high vs. low) and mutational frequency (high vs. low) (P < 0.05). **(E)** In never tobacco users, there was a significant positive correlation between TGEP scores and mutational frequency. This correlation was not present in current or former tobacco users. (* P < 0.05, ** P < 0.01)

TGEP scores significantly differed based on current, former, never tobacco use status (**Figure 5C**, P = 0.029), with current tobacco users having lower scores than never or former users (P = 0.028 and 0.085 respectively). Tobacco’s effect on the TIME was also evident when TGEP scores were analyzed in combination with tumor mutational frequency. We used previously published thresholds for TGEP (score greater than -0.318) and mutational frequency (greater than 86 somatic, nonsynonymous variants per sample) that have been previously shown to predict response to PD-1 inhibition in HNSCC alone or in combination with T cell inflamed GEP scores.^16^ These thresholds divided patients into four groups based on high and low TGEP score and mutational frequency values. Compared with former or never tobacco users, current tobacco users disproportionately distributed into groups where one or both of TGEP scores or mutational frequency were low (**Figure 5D**, P = 0.018). Never tobacco users demonstrated a significant positive correlation between TGEP score and mutational frequency (**Figure 5E**, Rho = 0.612, P = 0.006). This positive correlation was not present in former users (**Figure 5E**, Rho = 0.010, P = 0.974) and or current tobacco users (**Figure 5E**, Rho = -0.524, P = 0.197). Taken together, our results suggest that varying degrees of T cell-related mRNA and protein expression are present within HPV(+) OPSCC primary tumors, and that low expression of these markers is associated with tobacco use at time of diagnosis.

### T cell-inflamed TIME is associated with survival and treatment response

Similar to our unsupervised clustering results, associations between T cell inflammation and treatment outcomes were observed using TGEP scores. Patients with death and/or recurrence had lower TGEP scores compared to patients who survived without disease (P = 0.054). We performed cut point analysis, which revealed that at a TGEP score threshold of −0.235, higher TGEP scores were significantly associated with improved OS and DFS (**Figure 6A, B**; P = 0.002 and 0.012, respectively). IHC expression of CD3, CD4, CD8, and PD-L1 were all significantly greater in samples with GEP scores above this threshold (**Supplemental Figure S5B,D,F,H,** all FDR < 0.1).

**Figure 6:**
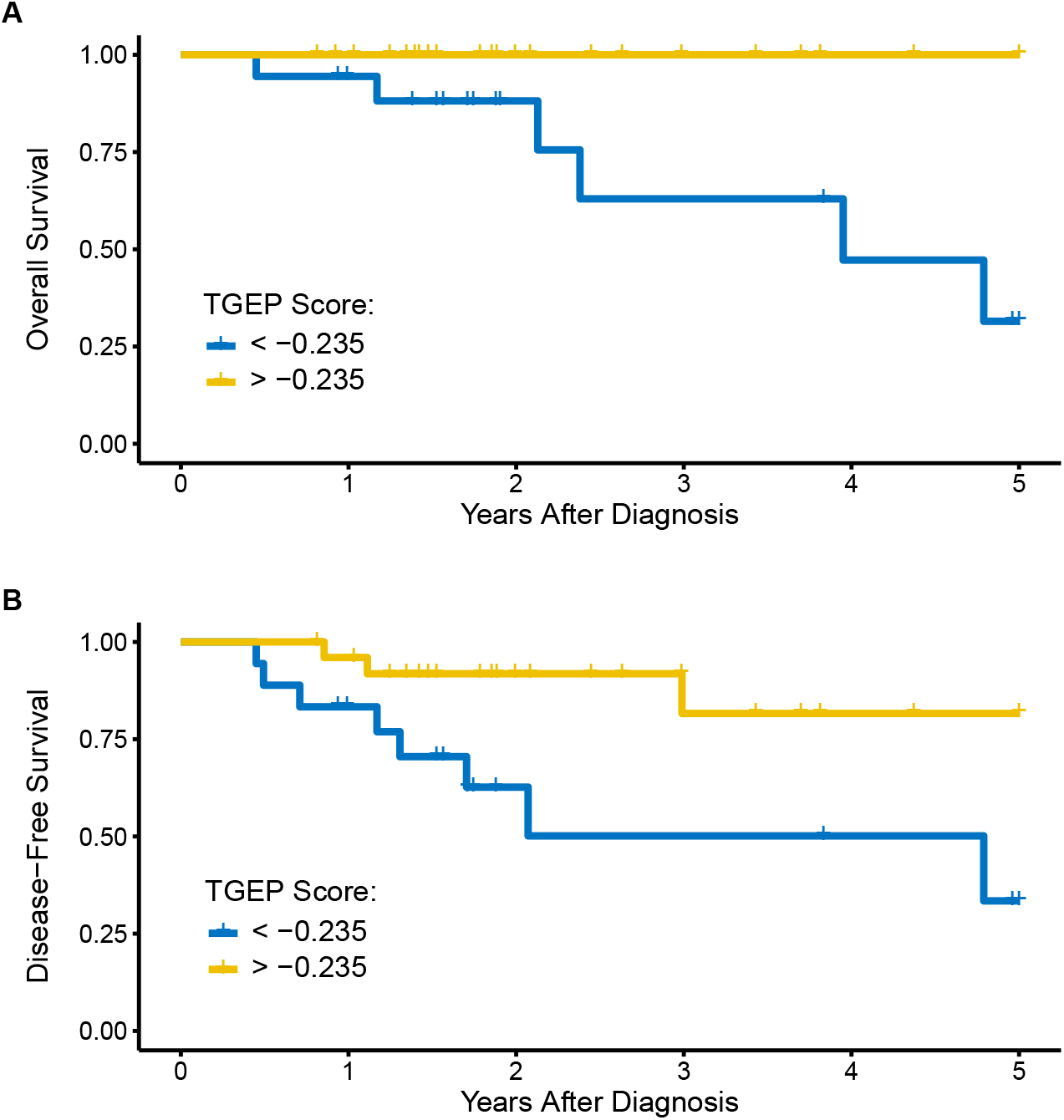
TGEP scores are significantly associated with oncologic outcomes. **(A)** OS was significantly reduced in patients with a TGEP < −0.235. (P < 0.01) **(B)** DFS was significantly reduced in patients with a TGEP < -0.235. (P < 0.05)

To ensure tissue specimen sources did not represent a source of bias in these results, we compared samples by primary treatment modality. When comparing primary surgery versus primary CRT, we did not identify differences in TGEP scores (**Supplemental Figure S5I,** P = 0.19, mean difference 0.13, 95% CI −0.07 to 0.33), nor did we identify categorical differences using the −0.235 TGEP score threshold (P = 0.31). We sought to determine if our observed trends related to tobacco and clinical outcomes were present in an independent cohort. We examined expression of the 18 transcripts in the TGEP score in p16-positive OPSCC patients from The Cancer Genome Atlas (TCGA) using normalized RNA sequencing data. In this cross-platform analysis, current tobacco users and patients with death and/or recurrence were similarly concentrated among patients with low expression of TGEP transcripts (**Supplemental Figure S6**).

## DISCUSSION

Using a multiomic approach, we have identified multiple biologic differences present in primary HPV(+) OPSCC tumors related to tobacco exposure. Most importantly, we demonstrate that current tobacco use is associated with decreased T cell infiltration of primary HPV(+) OPSCC tumors. Our WES results suggest this effect is present despite a lack of significant differences in TMB or recurrent oncogenic mutations associated with tobacco use. Further, low T cell infiltration was associated with decreased OS and DFS. Taken together, our results suggest that the primary and clinically relevant feature associated with tobacco use in HPV(+) OPSCC is decreased T cell infiltration of the primary TIME.

The immunosuppressive molecular effects of tobacco exposure have been previously demonstrated using a variety of experimental methods. In case-control studies and experimental work using human PBMC’s, tobacco exposure has been associated with cytokine alterations with particular relevance to both antiviral and antitumor immune responses, including decreases in IFN-y, IL-2, TNF, IL-15, and IL-16.^21–26^ Experimental evidence using animal models exposed to tobacco smoke also support this hypothesis, with multiple studies showing that tobacco exposure results in systemic T cell anergy and impaired immune responses to a variety of challenges, including viruses.^27–31^ Other investigations have revealed impairment of NK and dendritic cells in response to tobacco exposure.^32–35^ These studies provide biologic rationale for the association between tobacco and poor immune infiltrate identified in our study.

Tobacco-related immunosuppression has been previously noted in the context of human malignancies including HNSCC. Our results are consistent with a recent report demonstrating decreased CD8+ IHC staining in tobacco-exposed primary HPV(+) OPSCC tumors.^13^ Our investigation elucidates this phenomenon in much greater biologic detail by using a readout with multiple mRNA hybridization probes, validating in IHC, and controlling for mutational burden using our WES data. Multiple groups have described a similar inverse relationship between tobacco exposure and immune infiltration in HPV(-) HSNCC.^36–38^ The effect of tobacco may be site-specific in squamous cell carcinoma (SCC), as tobacco has been associated with an inflamed TIME in lung SCC.^38^ This difference in lung SCC may be related to tobacco-induced pulmonary inflammation which has been demonstrated outside of the context of cancer.^39^ Beyond SCC, tobacco has been associated with immunosuppression in primary colorectal carcinoma, another highly immunogenic tumor.^40,41^

Further, tobacco exposure influences HPV-specific immune responses. Multiple groups have demonstrated that tobacco use is associated with impaired immune responses and clearance of oncogenic HPV infections in human female genital infections.^42–44^ In the oropharynx specifically, current tobacco use is strongly associated with an increased incidence of oral HPV16 infections.^45^ These findings are important given the putative oncologic benefits of HPV in the context of HNSCC. HPV(+) OPSCC tumors display greater T cell infiltration compared to HPV(-) HNSCC, a finding that may contribute to superior oncologic outcomes in HPV(+) disease.^46–48^ The results of the present study are important given clinical evidence demonstrating that tobacco users with HPV(+) OPSCC have an attenuated oncologic prognosis compared to non-tobacco users.^2,7–9,49^ Based on our results, we speculate that this attenuated prognosis is related to tobacco-induced immunosuppression, which diminishes the immunologic benefit of having an HPV(+) tumor, forcing HPV(+) OPSCC tobacco users into an intermediate prognostic group in between tobacco-naїve HPV(+) OPSCC and HPV(-) OPSCC. Prospective studies are needed to precisely define the systemic and local immunologic features associated with tobacco use in HPV(+) OPSCC and how these features intersect with oncologic outcomes.

The TGEP score used in this study provided a useful method to quantitatively describe the spectrum of T cell-inflammation in our cohort. This score has been validated in large patient cohorts as a predictor of response to PD-1 inhibitors, is agnostic to tumor-type, and was developed on the same NanoString RNA hybridization platform used in this study.^15,16^ Its utility in stratifying patients by response to PD-1 inhibitors likely lies in its ability to identify which patients generated an initial antitumor immune response.^15^ The transcripts present in the TGEP characterize different components of this response: neoantigen presentation, chemoattraction and infiltration of cytotoxic T cells and professional antigen presenting cells, IFN-λ signaling, and the resulting upregulation of immune checkpoints.^15,50^ Like IHC staining, TGEP scores may be derived from formalin fixed paraffin embedded samples. However compared to IHC, TGEP scores are quantitative, operator-independent, and reliant on hybridization of multiple probes rather than the binding of a single antibody. Notably, overlap between low TGEP scores and tobacco use was incomplete, with a subset of former or never tobacco users also having low TGEP scores. These results suggest that current tobacco use is not required to achieve this high-risk phenotype. We propose that gene expression profiles like TGEP, which provide quantitative measures of TIME inflammation, may provide useful objective data for risk stratification in clinical trials aimed at HPV(+) OPSCC treatment deintensification and/or immunotherapeutics.

Our WES data also provide important insights related to tobacco exposure in HPV(+) OPSCC. Our results were consistent with the largest genomics study to date of HPV(+) OPSCC patients, which demonstrated that while mutational burden increased with heavy tobacco exposure in HPV(−) tumors, there was no difference in mutational burden in HPV(+) OPSCC on the basis of tobacco exposure.^12^ Similarly, we were unable to identify a higher rate of any particular mutation in tobacco users, including canonical HPV(-) mutations.^12^ We acknowledge that we were not powered to detect small effect sizes and that a much larger sequencing cohort of HPV(+) OPSCC patients will be required in order to identify more subtle differences in mutational profiles differ on the basis of tobacco use.

We did identify mutational differences related to tobacco use in our cohort in other analyses. We found that tobacco exposed tumors had higher rates of T>C substitutions, lower rates of APOBEC and aging-related mutational signatures, and a greater share of mutations that could not be classified into a COSMIC signature. Importantly, these effects were observed in both current and former tobacco users, whereas tobacco’s associations with the TIME were observed predominately in current tobacco users. The finding that our cohort totally lacked mutational signature 4, which is associated with tobacco use, is not unexpected. Compared to tobacco-naїve tumors, signature 4 is significantly increased in tobacco-exposed lung SCC, lung adenocarcinoma, and larynx SCC, but remains largely absent in other tobacco-associated tumors such as oral cavity, pharynx and esophageal SCC as well as bladder cancer.^51^ Among cancers without signature 4, significant increases T>C substitutions have been observed in tobacco exposed oral cavity and bladder tumors.^51^ Based on our findings and previous genomic studies, we conclude that while tobacco exposure in HPV(+) OPSCC is associated with discernible differences in somatic mutations, there remains no clear evidence that this affects mutational burden or recurrent oncogenic mutations. Although they have an intermediate prognosis, available evidence does not indicate that tobacco-exposed HPV(+) tumors represent a mutational intermediary between tobacco naїve HPV(+) OPSCC and HPV(-) OPSCC.

We identified multiple genes that have yet to be described as significantly mutated in HPV(+) OPSCC. *B2M* variants were previously noted for HNSCC and HPV(+) OPSCC specifically in TCGA, although criteria for statistical significance were not met^18^. B2M is a component of the MHC-I complex and serves an indispensable role in antigen presentation.^52^ Another identified novel mutated gene with potential biologic relevance was *SMARCAL1*, which codes for an annealing helicase that limits genomic damage at stalled replication forks.^53^ Like p53, SMARCAL1 is also phosphorylated by ATM serine/threonine kinase, whose gene is deleted in recurrent 11q22 copy number losses in HPV(+) OPSCC.^18^ Further work is needed to clarify the role of SMARCAL1 relative to other known sources of genomic instability in HPV(+) OPSCC. The biologic relevance of the other identified mutations *AK5, IFI27, IQGC,* and *METTL24* in HPV(+) OPSCC is not clear and will also require further investigation.

The present study has several limitations not previously mentioned. Our study’s most significant limitation was that we relied on retrospective review of the medical record to determine tobacco exposure. While we confidently classified all patients into current, former, or never tobacco status in all patients, determining pack years was not possible in many cases. This limitation may be avoided in future studies through administration of a tobacco survey at the time of enrollment. Even with these measures in place, biases associated with patient-reported tobacco use are never entirely avoided, a fact that underscores the importance of identifying objective biomarkers of high-risk disease that do not rely on patient-reported behavioral data. We also acknowledge that while mRNA hybridization provides a platform that is more readily converted to a clinical assay, its utility as a tool for discovery is limited compared to RNA sequencing. Although the TGEP score we used has had extensive prior validation and correlated well with IHC staining in our cohort, other experimental approaches will be needed to better define the mechanisms associated with smoking-related immunosuppression in HPV(+) OPSCC.

In summary, we have shown that current tobacco use is associated with decreased immune infiltration in HPV(+) OPSCC. Although we identified tobacco-associated differences in single base substitutions and mutational signatures, tobacco was not associated with increased TMB or differences in recurrent oncogenic mutations. TGEP scores are associated with OS and DFS in HPV(+) OPSCC and may represent a quantitative clinical assay for use in future risk stratification efforts. We conclude that in HPV(+) OPSCC, the primary and clinically relevant association with tobacco use is decreased T cell infiltration of the TIME. This works suggests that immunosuppression may account for the increased oncologic risk observed in tobacco users with HPV(+) OPSCC.

## REFERENCES

1. Chaturvedi, A. K. et al. Human papillomavirus and rising oropharyngeal cancer incidence in the United States. J. Clin. Oncol. 29, 4294–4301 (2011).

2. Ang, K. K. et al. Human papillomavirus and survival of patients with oropharyngeal cancer. N. Engl. J. Med. 363, 24–35 (2010).

3. Centers for Disease Control and Prevention. Cancers Associated with Human Papillomavirus, United States—2012-2016. USCS Data Brief, no 10 (2019).

4. Sinha, P. et al. Survival for HPV-positive oropharyngeal squamous cell carcinoma with surgical versus non-surgical treatment approach: A systematic review and meta-analysis. Oral Oncol. 86, 121–131 (2018).

5. Wahle, B. & Zevallos, J. Transoral Robotic Surgery and De-escalation of Cancer Treatment. Otolaryngologic clinics of North America (2020) doi:10.1016/j.otc.2020.07.009.

6. Gillison, M. L. et al. Tobacco smoking and increased risk of death and progression for patients with p16-positive and p16-negative oropharyngeal cancer. J. Clin. Oncol. 30, 2102–2111 (2012).

7. Hafkamp, H. C. et al. Marked differences in survival rate between smokers and nonsmokers with HPV 16-associated tonsillar carcinomas. Int. J. Cancer 122, 2656–2664 (2008).

8. Lin, B. M. et al. Long-term prognosis and risk factors among patients with HPV-associated oropharyngeal squamous cell carcinoma. Cancer 119, 3462–3471 (2013).

9. Fakhry, C. et al. Human Papillomavirus and Overall Survival After Progression of Oropharyngeal Squamous Cell Carcinoma. J Clin Oncol 32, 3365–3373 (2014).

10. Elhalawani, H. et al. Tobacco exposure as a major modifier of oncologic outcomes in human papillomavirus (HPV) associated oropharyngeal squamous cell carcinoma. BMC Cancer 20, (2020).

11. Maxwell, J. H. et al. Tobacco use in HPV-positive advanced oropharynx cancer patients related to increased risk of distant metastases and tumor recurrence. Clin Cancer Res 16, 1226 (2010).

12. Gillison, M. L. et al. Human papillomavirus and the landscape of secondary genetic alterations in oral cancers. Genome Res. (2018) doi:10.1101/gr.241141.118.

13. Kemnade, J. O. et al. CD8 infiltration is associated with disease control and tobacco exposure in intermediate-risk oropharyngeal cancer. Sci Rep 10, 243 (2020).

14. Liao, Y., Wang, J., Jaehnig, E. J., Shi, Z. & Zhang, B. WebGestalt 2019: gene set analysis toolkit with revamped UIs and APIs. Nucleic Acids Research 47, W199–W205 (2019).

15. Ayers, M. et al. IFN-γ-related mRNA profile predicts clinical response to PD-1 blockade. J Clin Invest 127, 2930–2940 (2017).

16. Cristescu, R. et al. Pan-tumor genomic biomarkers for PD-1 checkpoint blockade–based immunotherapy. Science 362, (2018).

17. Amin, D. et al. Metformin Effects on FOXP3+ and CD8+ T Cell Infiltrates of Head and Neck Squamous Cell Carcinoma. Laryngoscope 130, E490–E498 (2020).

18. Lawrence, M. S. et al. Comprehensive genomic characterization of head and neck squamous cell carcinomas. Nature 517, 576–582 (2015).

19. Cho, A. et al. MUFFINN: cancer gene discovery via network analysis of somatic mutation data. Genome Biology 17, 129 (2016).

20. Alexandrov, L. B. et al. Signatures of mutational processes in human cancer. Nature 500, 415–421 (2013).

21. Shiels, M. S. et al. Cigarette Smoking and Variations in Systemic Immune and Inflammation Markers. J Natl Cancer Inst 106, (2014).

22. Mian, M. F., Stämpfli, M. R., Mossman, K. L. & Ashkar, A. A. Cigarette smoke attenuation of poly I:C-induced innate antiviral responses in human PBMC is mainly due to inhibition of IFN-beta production. Mol Immunol 46, 821–829 (2009).

23. Mian, M. F., Pek, E. A., Mossman, K. L., Stämpfli, M. R. & Ashkar, A. A. Exposure to cigarette smoke suppresses IL-15 generation and its regulatory NK cell functions in poly I:C-augmented human PBMCs. Mol Immunol 46, 3108–3116 (2009).

24. Liu, G., Arimilli, S., Savage, E. & Prasad, G. L. Cigarette smoke preparations, not moist snuff, impair expression of genes involved in immune signaling and cytolytic functions. Sci Rep 9 (2019).

25. Ma, M., Lj, M., S, E.-M. & Dc, L. Inhibition of IFN-gamma-dependent antiviral airway epithelial defense by cigarette smoke. Respiratory research vol. 11 (2010).

26. van Dijk, A. P. et al. Transdermal nicotine inhibits interleukin 2 synthesis by mononuclear cells derived from healthy volunteers. Eur J Clin Invest 28, 664–671 (1998).

27. Geng, Y., Savage, S. M., Johnson, L. J., Seagrave, J. & Sopori, M. L. Effects of nicotine on the immune response. I. Chronic exposure to nicotine impairs antigen receptor-mediated signal transduction in lymphocytes. Toxicol Appl Pharmacol 135, 268–278 (1995).

28. Geng, Y., Savage, S. M., Razani-Boroujerdi, S. & Sopori, M. L. Effects of nicotine on the immune response. II. Chronic nicotine treatment induces T cell anergy. The Journal of Immunology 156, 2384–2390 (1996).

29. Kalra, R., Singh, S. P., Savage, S. M., Finch, G. L. & Sopori, M. L. Effects of cigarette smoke on immune response: chronic exposure to cigarette smoke impairs antigen-mediated signaling in T cells and depletes IP3-sensitive Ca(2+) stores. J Pharmacol Exp Ther 293, 166–171 (2000).

30. Feng, Y. et al. Exposure to cigarette smoke inhibits the pulmonary T-cell response to influenza virus and Mycobacterium tuberculosis. Infect Immun 79, 229–237 (2011).

31. Schierl, M. et al. Tobacco Smoke-Induced Immunologic Changes May Contribute to Oral Carcinogenesis. J Investig Med 62, 316–323 (2014).

32. Hao, J. et al. Nicotinic receptor β2 determines NK cell-dependent metastasis in a murine model of metastatic lung cancer. PLoS One 8, e57495 (2013).

33. Lu, L.-M. et al. Cigarette smoke impairs NK cell-dependent tumor immune surveillance. J Immunol 178, 936–943 (2007).

34. Alkhattabi, N., Todd, I., Negm, O., Tighe, P. J. & Fairclough, L. C. Tobacco smoke and nicotine suppress expression of activating signaling molecules in human dendritic cells. Toxicol Lett 299, 40–46 (2018).

35. Vassallo, R., Tamada, K., Lau, J. S., Kroening, P. R. & Chen, L. Cigarette smoke extract suppresses human dendritic cell function leading to preferential induction of Th-2 priming. J Immunol 175, 2684–2691 (2005).

36. Foy, J.-P. et al. The immune microenvironment of HPV-negative oral squamous cell carcinoma from never-smokers and never-drinkers patients suggests higher clinical benefit of IDO1 and PD1/PD-L1 blockade. Ann Oncol 28, 1934–1941 (2017).

37. Iglesia, J. V. de la et al. Effects of Tobacco Smoking on the Tumor Immune Microenvironment in Head and Neck Squamous Cell Carcinoma. Clin Cancer Res 26, 1474–1485 (2020).

38. Desrichard, A. et al. Tobacco Smoking-Associated Alterations in the Immune Microenvironment of Squamous Cell Carcinomas. J Natl Cancer Inst 110, 1386–1392 (2018).

39. Hodge, G., Nairn, J., Holmes, M., Reynolds, P. N. & Hodge, S. Increased intracellular T helper 1 proinflammatory cytokine production in peripheral blood, bronchoalveolar lavage and intraepithelial T cells of COPD subjects. Clin Exp Immunol 150, 22–29 (2007).

40. Hamada, T. et al. Smoking and Risk of Colorectal Cancer Sub-Classified by Tumor-Infiltrating T Cells. J Natl Cancer Inst 111, 42–51 (2018).

41. Fujiyoshi, K. et al. Smoking Status at Diagnosis and Colorectal Cancer Prognosis According to Tumor Lymphocytic Reaction. JNCI Cancer Spectr 4, (2020).

42. Simen-Kapeu, A. et al. Smoking impairs human papillomavirus (HPV) type 16 and 18 capsids antibody response following natural HPV infection. Scand J Infect Dis 40, 745–751 (2008).

43. Giuliano, A. R. et al. Clearance of oncogenic human papillomavirus (HPV) infection: effect of smoking (United States). Cancer Causes Control 13, 839–846 (2002).

44. Koshiol, J. et al. Smoking and time to clearance of human papillomavirus infection in HIV-seropositive and HIV-seronegative women. Am J Epidemiol 164, 176–183 (2006).

45. Fakhry, C., Gillison, M. L. & D’Souza, G. Tobacco use and oral HPV16 infection. JAMA 312, 1465–1467 (2014).

46. Mandal, R. et al. The head and neck cancer immune landscape and its immunotherapeutic implications. JCI Insight 1,.

47. Jung, A. C. et al. CD8-alpha T-cell infiltration in human papillomavirus-related oropharyngeal carcinoma correlates with improved patient prognosis. International Journal of Cancer 132, E26–E36 (2013).

48. Masterson, L. et al. CD8+ T cell response to human papillomavirus 16 E7 is able to predict survival outcome in oropharyngeal cancer. Eur. J. Cancer67, 141–151 (2016).

49. Chen, S. Y. et al. The association of smoking and outcomes in HPV-positive oropharyngeal cancer: A systematic review. Am J Otolaryngol 41, 102592 (2020).

50. Spranger, S. & Gajewski, T. F. Impact of oncogenic pathways on evasion of antitumour immune responses. Nat. Rev. Cancer 18, 139–147 (2018).

51. Alexandrov, L. B. et al. Mutational signatures associated with tobacco smoking in human cancer. Science 354, 618–622 (2016).

52. D’Urso, C. M. et al. Lack of HLA class I antigen expression by cultured melanoma cells FO-1 due to a defect in B2m gene expression. J. Clin. Invest. 87, 284–292 (1991).

53. Bansbach, C. E., Bétous, R., Lovejoy, C. A., Glick, G. G. & Cortez, D. The annealing helicase SMARCAL1 maintains genome integrity at stalled replication forks. Genes Dev 23, 2405–2414 (2009).

