## Supplemental Figures for "Comparative Multiomic Analysis Reveals Low T Cell Infiltration as the Primary Feature of Tobacco Use in HPV(+) Oropharyngeal Cancer"

#### Figure S1

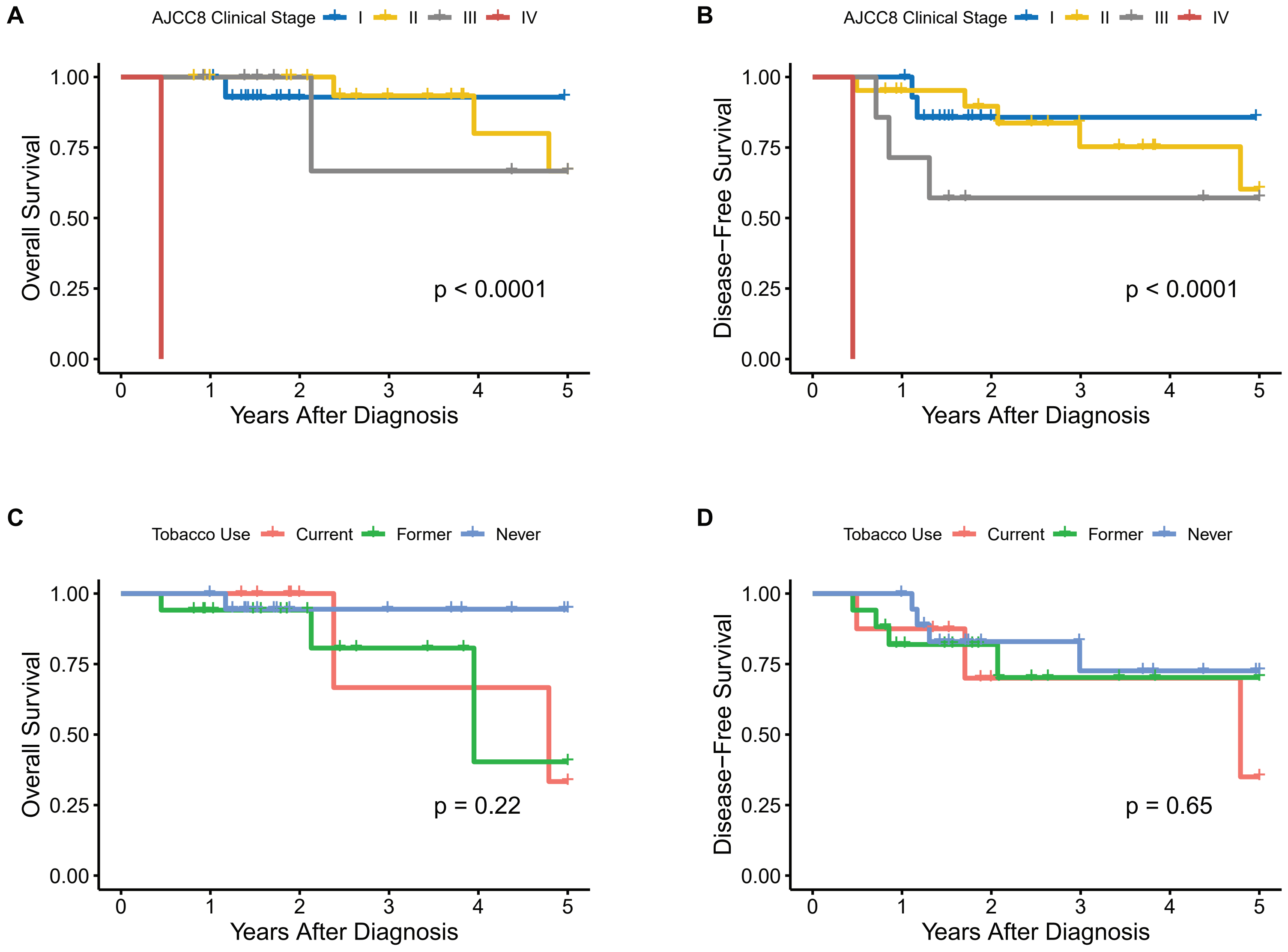
A
AJCC8 Clinical Stage I II III IV
Overall Survival
1.00
0.75
0.50
0.25
0.00
p < 0.0001
0 1 2 3 4 5
Years After Diagnosis
B
AJCC8 Clinical Stage I II III IV
Disease−Free Survival
p < 0.0001
Years After Diagnosis
C
Tobacco Use Current Former Never
Overall Survival
p = 0.22
Years After Diagnosis
D
Tobacco Use Current Former Never
Disease−Free Survival
p = 0.65
Years After Diagnosis

### Figure S2

A

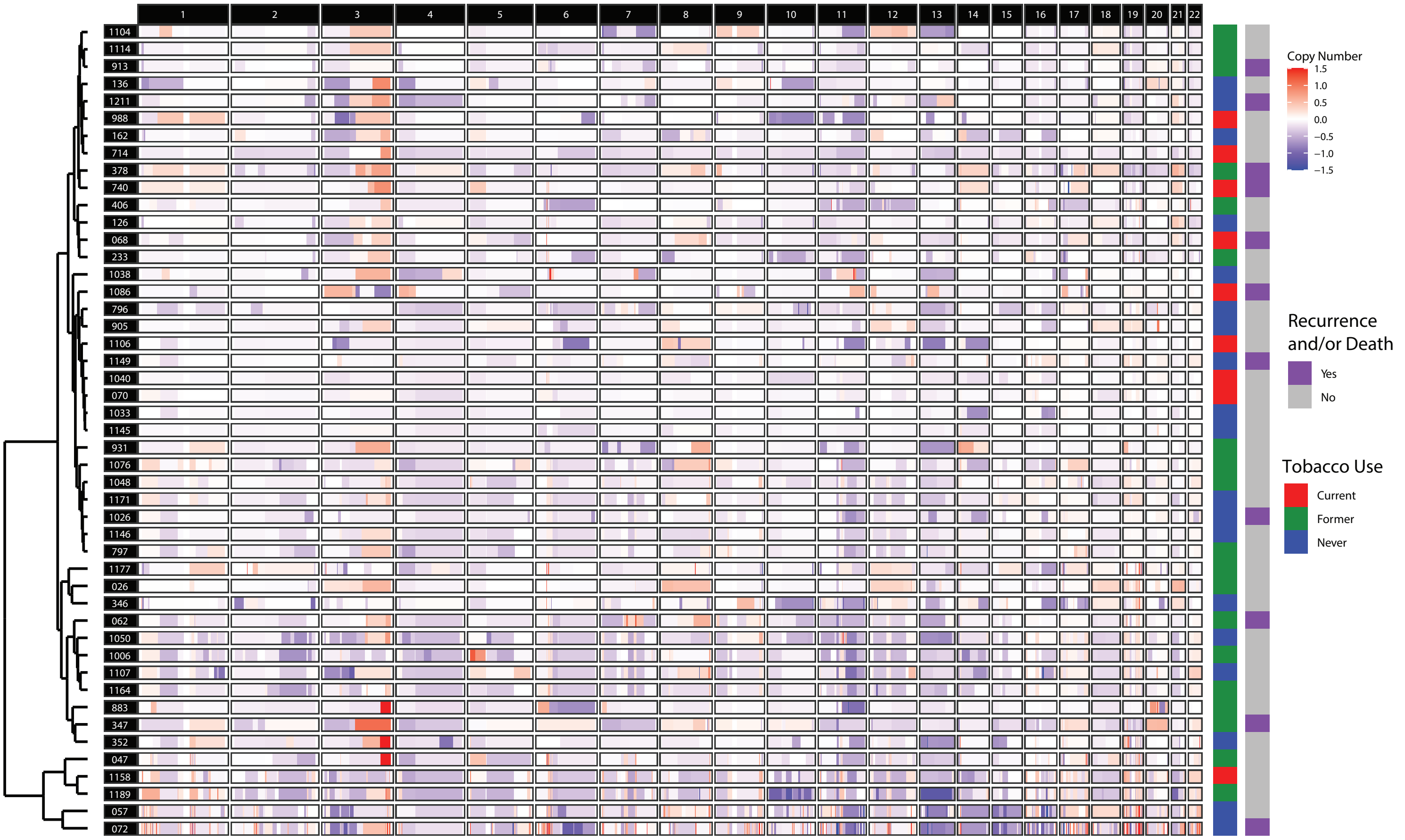

1 2 3 4 5 6 7 8 9 10 11 12 13 14 15 16 17 18 19 20 21 22
1104
1114
913
136
1211
988
162
714
378
740
406
126
068
233
1038
1086
796
905
1106
1149
1040
070
1033
1145
931
1076
1048
1171
1026
1146
797
1177
026
346
062
1050
1006
1107
1164
883
347
352
047
1158
1189
057
072
Copy Number
1.5
1.0
0.5
0.0
−0.5
−1.0
−1.5
Recurrence and/or Death
Yes
No
Tobacco Use
Current
Former
Never


B

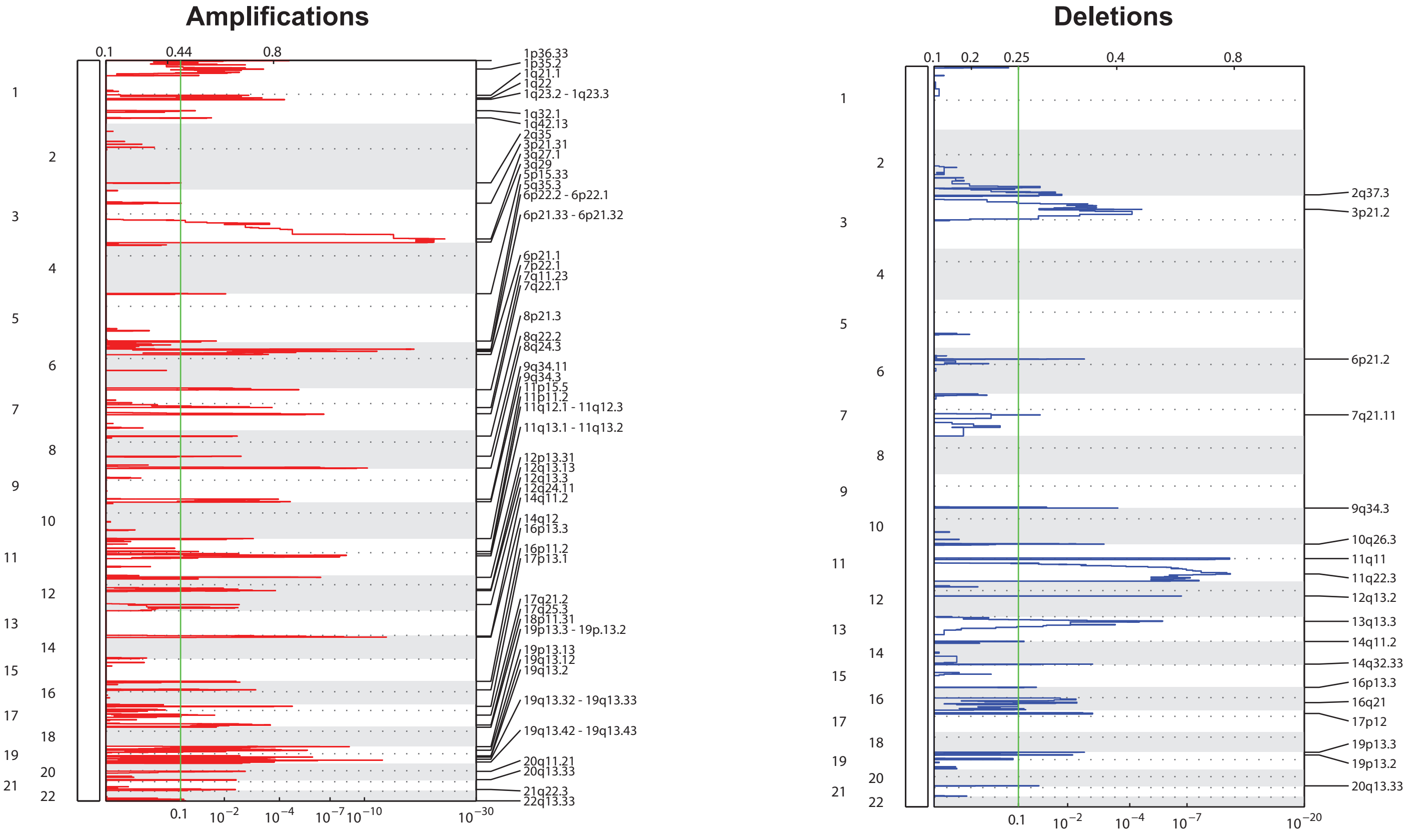

Amplifications
0.1
0.44
0.8
1p36.33
1p35.2
1q21.1
1q22
1q23.2 - 1q23.3
1q32.1
1q42.13
2q35
3p21.31
3q27.1
3q29
5p15.33
5q35.3
6p22.2 - 6p22.1
6p21.33 - 6p21.32
6p21.1
7p22.1
7q11.23
7q22.1
8p21.3
8q22.2
8q24.3
9q34.11
9q34.3
11p15.5
11p11.2
11q12.1 - 11q12.3
11q13.1 - 11q13.2
12p13.31
12q13.13
12q13.3
12q24.11
14q11.2
14q12
16p13.3
16p11.2
17p13.1
17q21.2
17q25.3
18p11.31
19p13.3 - 19p.13.2
19p13.13
19q13.12
19q13.2
19q13.32 - 19q13.33
19q13.42 - 19q13.43
20q11.21
20q13.33
21q22.3
22q13.33
0.1
$10^{-2}$
$10^{-4}$
$10^{-7}$
$10^{-10}$
$10^{-30}$
Deletions
0.1
0.2
0.25
0.4
0.8
2q37.3
3p21.2
6p21.2
7q21.11
9q34.3
10q26.3
11q11
11q22.3
12q13.2
13q13.3
14q11.2
14q32.33
16p13.3
16q21
17p12
19p13.3
19p13.2
20q13.33
0.1
$10^{-2}$
$10^{-4}$
$10^{-7}$
$10^{-20}$

#### Figure S3

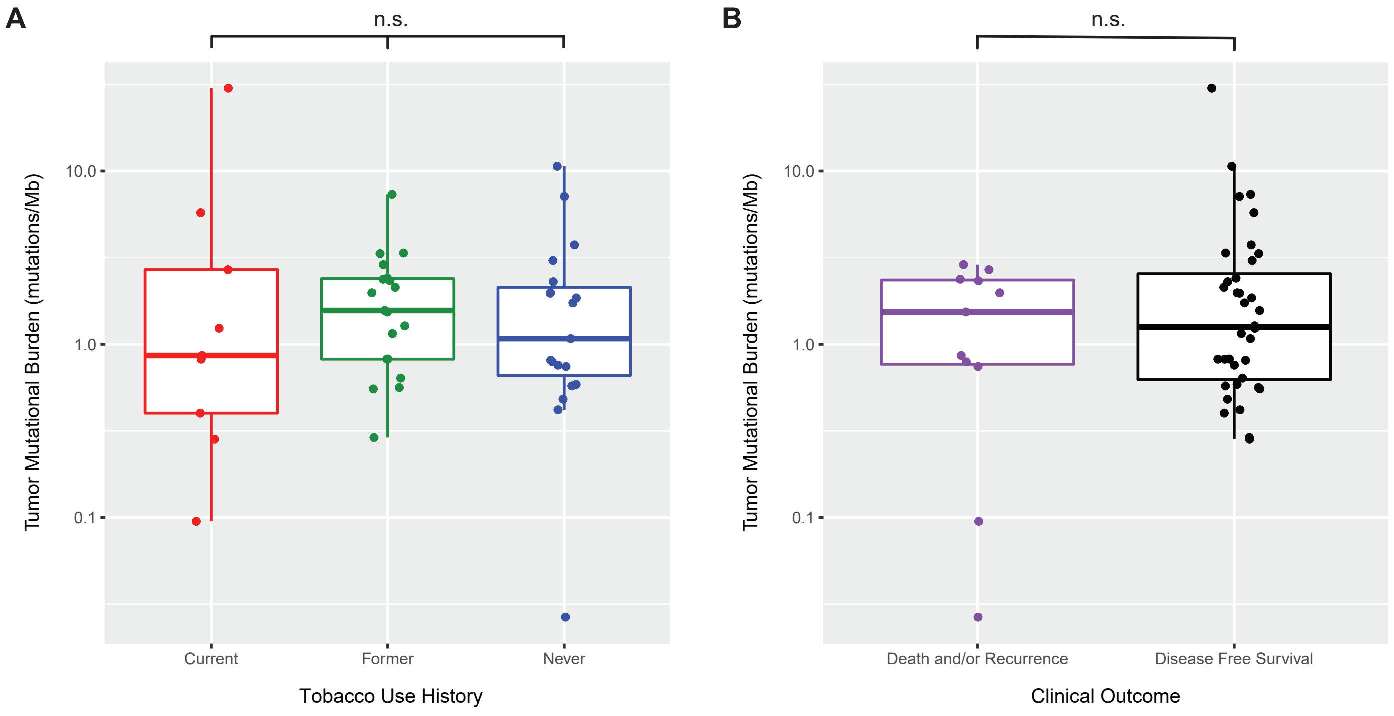
A
n.s.
Tumor Mutational Burden (mutations/Mb)
10.0
1.0
0.1
Current
Former
Never
Tobacco Use History
B
n.s.
Tumor Mutational Burden (mutations/Mb)
10.0
1.0
0.1
Death and/or Recurrence
Disease Free Survival
Clinical Outcome

**Figure S4**

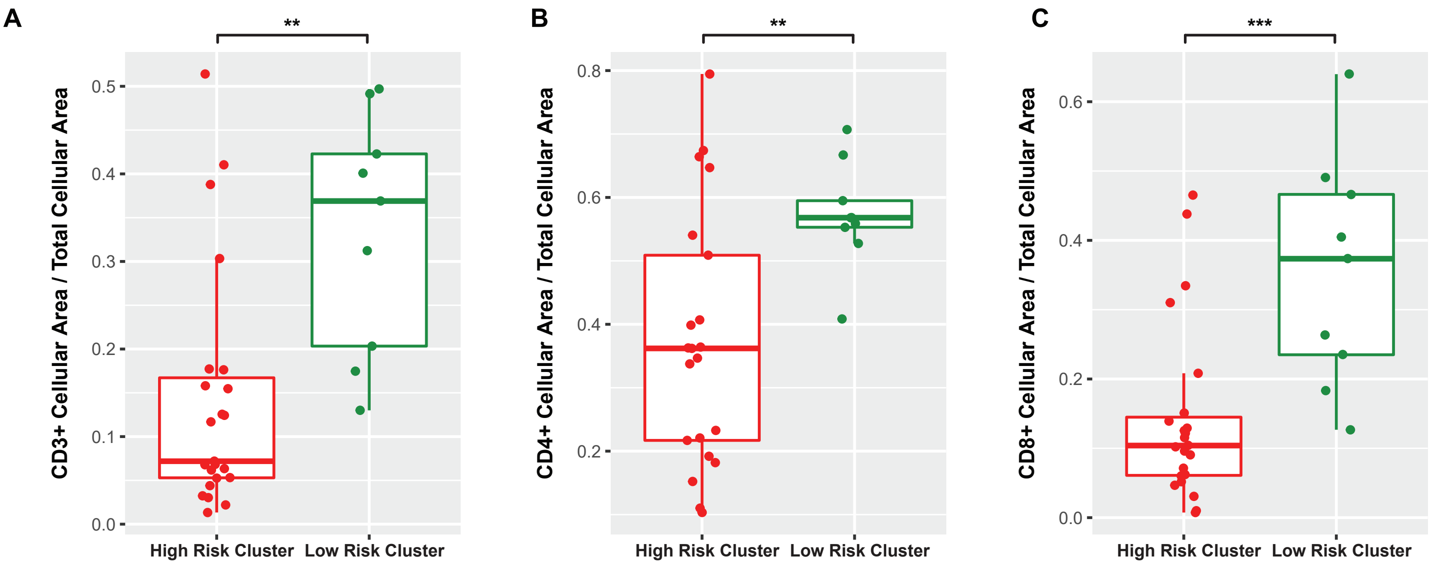
A
**
CD3+ Cellular Area / Total Cellular Area
0.5
0.4
0.3
0.2
0.1
0.0
High Risk Cluster
Low Risk Cluster
B
**
CD4+ Cellular Area / Total Cellular Area
0.8
0.6
0.4
0.2
High Risk Cluster
Low Risk Cluster
C
***
CD8+ Cellular Area / Total Cellular Area
0.6
0.4
0.2
0.0
High Risk Cluster
Low Risk Cluster

### Figure S5

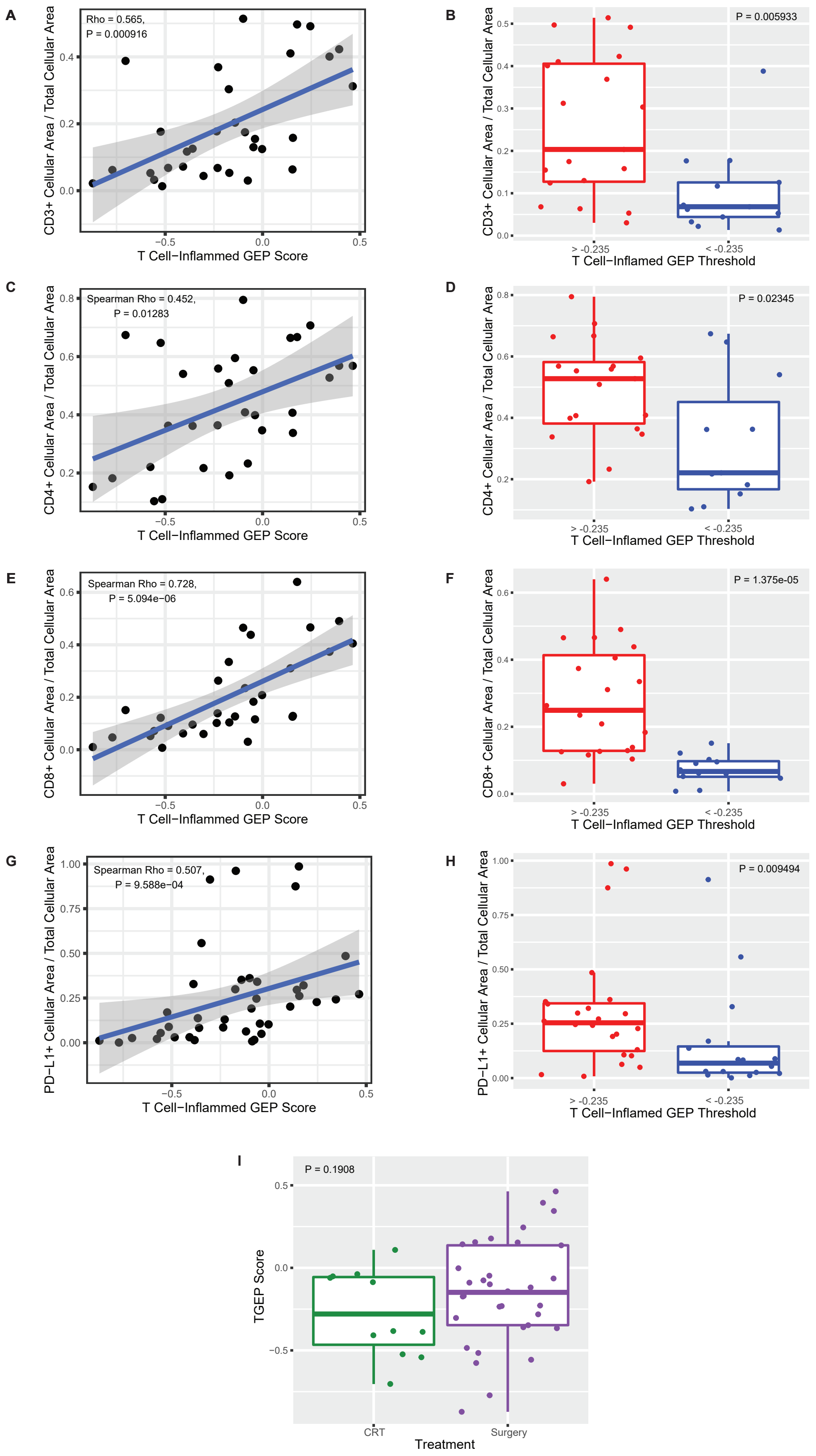

A
Rho = 0.565,
P = 0.000916
CD3+ Cellular Area / Total Cellular Area
T Cell−Inflammed GEP Score
B
P = 0.005933
CD3+ Cellular Area / Total Cellular Area
> -0.235
< -0.235
T Cell−Inflamed GEP Threshold
C
Spearman Rho = 0.452,
P = 0.01283
CD4+ Cellular Area / Total Cellular Area
T Cell−Inflammed GEP Score
D
P = 0.02345
CD4+ Cellular Area / Total Cellular Area
> -0.235
< -0.235
T Cell−Inflamed GEP Threshold
E
Spearman Rho = 0.728,
P = 5.094e−06
CD8+ Cellular Area / Total Cellular Area
T Cell−Inflammed GEP Score
F
P = 1.375e-05
CD8+ Cellular Area / Total Cellular Area
> -0.235
< -0.235
T Cell−Inflamed GEP Threshold
G
Spearman Rho = 0.507,
P = 9.588e−04
PD−L1+ Cellular Area / Total Cellular Area
T Cell−Inflammed GEP Score
H
P = 0.009494
PD−L1+ Cellular Area / Total Cellular Area
> -0.235
< -0.235
T Cell−Inflamed GEP Threshold
I
P = 0.1908
TGEP Score
CRT
Surgery
Treatment

### Figure S6

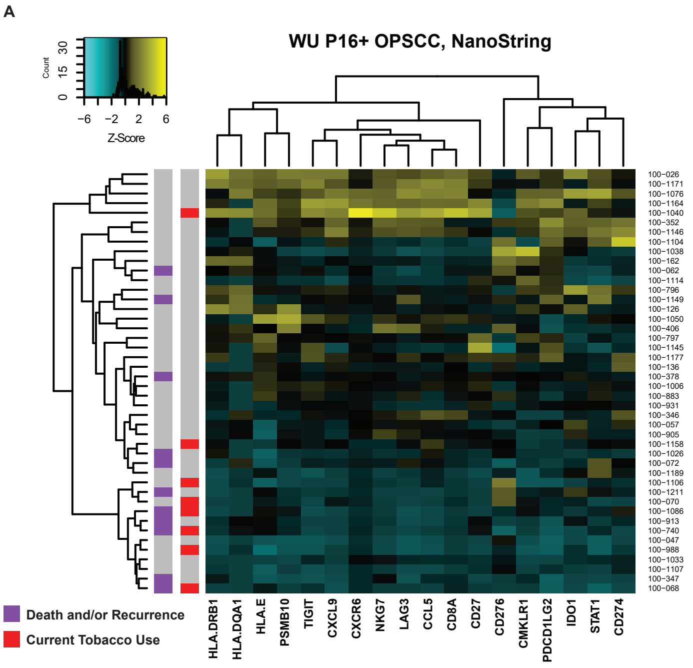
A
WU P16+ OPSCC, NanoString
Count
0
15
30
−6
−2
2
6
Z-Score
100−026
100−1171
100−1076
100−1164
100−1040
100−352
100−1146
100−1104
100−1038
100−162
100−062
100−1114
100−796
100−1149
100−126
100−1050
100−406
100−797
100−1145
100−1177
100−136
100−378
100−1006
100−883
100−931
100−346
100−057
100−905
100−1158
100−1026
100−072
100−1189
100−1106
100−1211
100−070
100−1086
100−913
100−740
100−047
100−988
100−1033
100−1107
100−347
100−068
HLA.DRB1
HLA.DQA1
HLA.E
PSMB10
TIGIT
CXCL9
CXCR6
NKG7
LAG3
CCL5
CD8A
CD27
CD276
CMKLR1
PDCD1LG2
IDO1
STAT1
CD274
Death and/or Recurrence
Current Tobacco Use

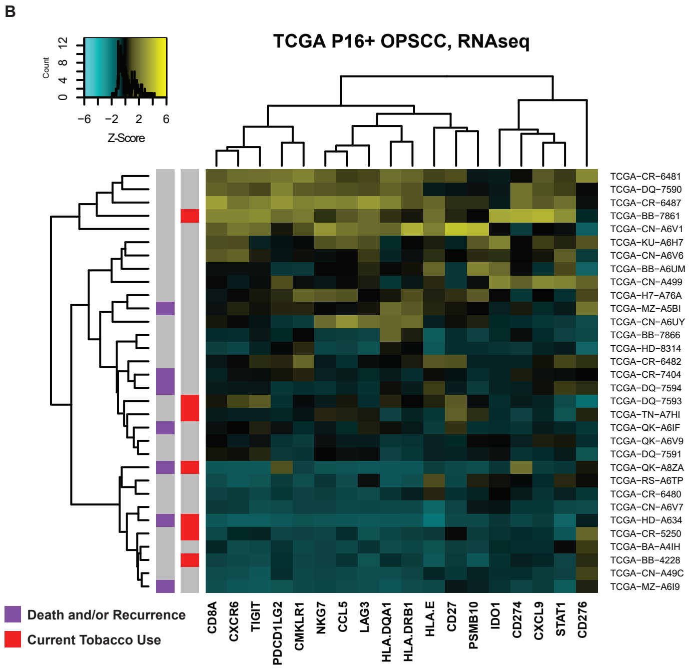
B
TCGA P16+ OPSCC, RNAseq
Count
0
4
8
12
−6
−2
2
6
Z-Score
TCGA−CR−6481
TCGA−DQ−7590
TCGA−CR−6487
TCGA−BB−7861
TCGA−CN−A6V1
TCGA−KU−A6H7
TCGA−CN−A6V6
TCGA−BB−A6UM
TCGA−CN−A499
TCGA−H7−A76A
TCGA−MZ−A5BI
TCGA−CN−A6UY
TCGA−BB−7866
TCGA−HD−8314
TCGA−CR−6482
TCGA−CR−7404
TCGA−DQ−7594
TCGA−DQ−7593
TCGA−TN−A7HI
TCGA−QK−A6IF
TCGA−QK−A6V9
TCGA−DQ−7591
TCGA−QK−A8ZA
TCGA−RS−A6TP
TCGA−CR−6480
TCGA−CN−A6V7
TCGA−HD−A634
TCGA−CR−5250
TCGA−BA−A4IH
TCGA−BB−4228
TCGA−CN−A49C
TCGA−MZ−A6I9
CD8A
CXCR6
TIGIT
PDCD1LG2
CMKLR1
NKG7
CCL5
LAG3
HLA.DQA1
HLA.DRB1
HLA.E
CD27
PSMB10
IDO1
CD274
CXCL9
STAT1
CD276
Death and/or Recurrence
Current Tobacco Use

#### Figure S7

CD8+ with hematoxylin

Scored images after RBG filters

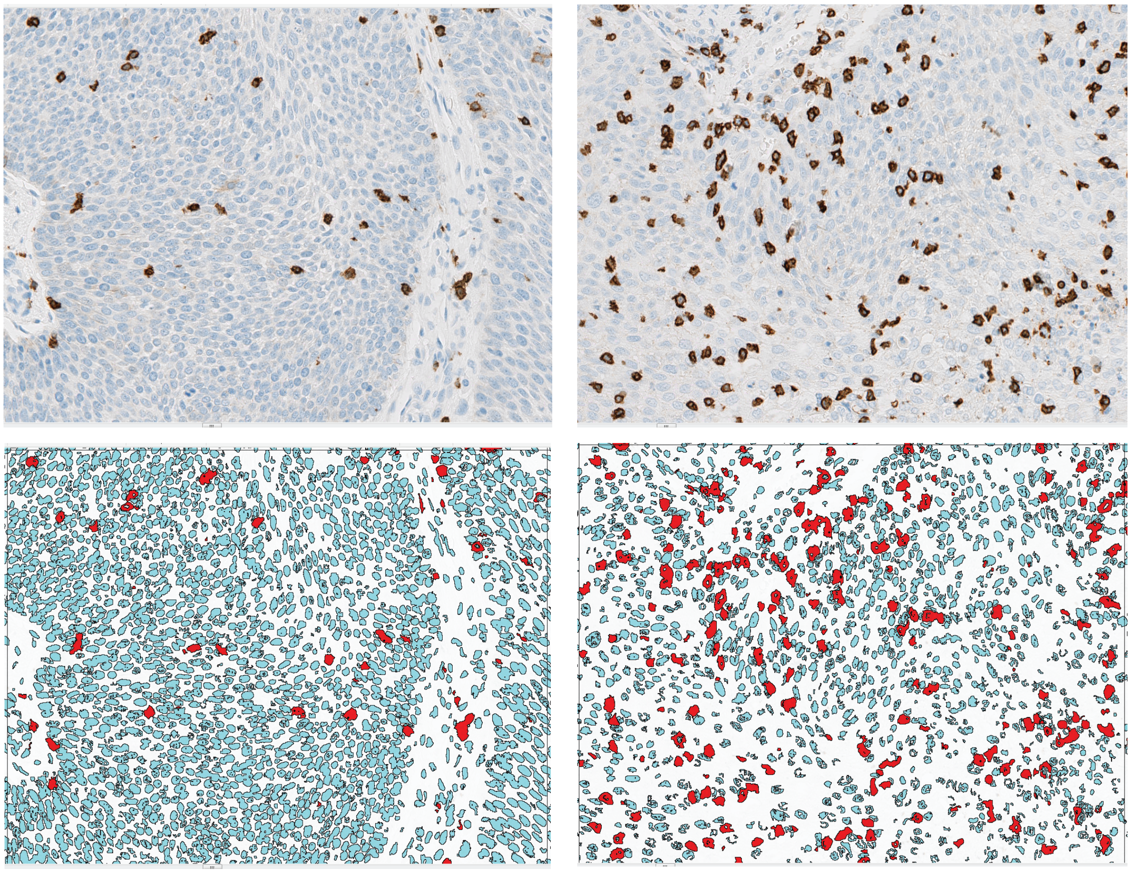

#### SUPPLEMENTAL FIGURE LEGENDS

**Figure S1**: Survival analyses for disease stage and tobacco use. Significant differences in **(A)** OS and **(B)** DFS were present by AJCC 8th Edition Clinical Stage. No significant differences in **(C)** OS and **(D)** DFS were present by tobacco use history.

**Figure S2:** Copy number alterations in HPV(+) OPSCC. **(A)** Heatmap displaying copy number alterations for each sample (rows) and chromosome (columns). Clinical annotation is included on the left margin. **(B)** GISTIC plots for focal amplifications and deletions in our cohort.

**Figure S3:** No significant differences in tumor mutational burden were present on the basis of **(A)** tobacco use history or **(B)** disease-free survival at five years. (ns = not significant)

**Figure S4** IHC confirmation of unsupervised clustering results. ORA revealed that gene cluster 4 was significantly enriched for adaptive immune-related pathways. IHC confirmed that the low-risk patient cluster, which had high expression of gene cluster 4, had significantly greater infiltration of **(A)** CD3+, **(B)** CD4+ and **(C)** CD8+ cells. (** $P < 0.01$, *** $P < 0.001$, all $FDR < 0.1$).

**Figure S5:** IHC validation of TGEP scores. **(A, C, E, G)** TGEP scores were significantly correlated with IHC markers for T cells and PD-L1+ cells. (all $P < 0.05$ and $FDR < 0.1$) **(B, D, F, H)** The TGEP cutpoint of -0.235, which was associated with significant differences in OS and DFS in our cohort (**Figure 5**), was also associated with significant differences in staining for T cell markers and PD-L1+ cells. (all $P < 0.05$ and $FDR < 0.1$) **(I)** No significant differences in TGEP scores were present on the basis of primary treatment modality (primary chemoradiation versus surgery, $P = 0.191$), suggesting that tissue sourcing (biopsy versus primary surgical specimen) does not represent a source of bias.

**Figure S6:** Cross-platform comparison of TGEP transcript expression. Scaled transcript expression was clustered in **(A)** the study cohort and **(B)** p16(+) OPSCC samples from TCGA, then annotated with current tobacco status and death and/or recurrence.

**Figure S7:** Representative images of high-resolution digital analysis of IHC staining. Custom red-green-blue (RBG) color filters were programmed which reliably distinguished and quantified areas of 3,3'-diaminobenzidine stained cells (red), hematoxylin stained cells (blue), and acellular background (white) over the entirety of a sample. Example images show original images at 20x magnification (top panels) and the same regions after application of filters (bottom panels).
