## Supplemental Methods for "Comparative Multiomic Analysis Reveals Low T Cell Infiltration as the Primary Feature of Tobacco Use in HPV(+) Oropharyngeal Cancer"

### Sequence Alignment, Somatic Variant Calling, Filtering, and Significance

Exome sequencing data was processed with the February 2019 version of the common workflow language somatic_exome pipeline developed at the McDonnell Genome Institute (https://github.com/genome/analysis-workflows). Pipeline execution was performed using Cromwell. Tracking of sample metadata and analysis results was performed using the McDonnell Genome Institute's Genome Modeling System.[1] Briefly, reads were aligned with bwa-mem (0.7.15) to version GRCh38 of the human genome reference supplemented with HLA decoy sequences (ftp://ftp.1000genomes.ebi.ac.uk/vol1/ftp/technical/reference/GRCh38_reference_genome/). Duplicate reads were marked with picard (2.18.1) and a base quality score recalibration applied with GATK (3.6).[2] Variant calling was performed using the union of calls from Strelka (2.9.9),[3] MuTect (3.6),[4] VarScan (2.4.2)[5] and Pindel (0.2.5b8)[6] and variants were annotated with the Variant Effect Predictor (Ensembl 93)[7]. Read counting of variants was performed via bam-readcount (0.7).

In order to obtain a final variant list, pipeline variants were further refined as follows. The variant list was filtered in R such that a candidate variant must have had > 4 variant supporting reads in the tumor sample, tumor variant allele frequency > 0.05, normal read depth > 20, and normal variant allele frequency <= 0.05. Non-coding and synonymous variants were removed from the candidate variant list. Using features described in deepSVR[8], variants were then classified as “Somatic” or “Failed” using an Extreme Gradient Boosting classifier via the XGB (booster = "gbtree", eta = 0.1, min_child_weight=1, max_depth = 8, gamma=0, subsample=1, scale_pos_weight=1, eval_metric = "auc") R library with a binary logistic classification objective. The classifier was trained on a subset of manually reviewed variants from the cohort. Five-fold cross validation of the model yielded an AUC of 0.85. Variants with a binary logistic probability between 0.35 and 0.65 were further refined via manual review.[9] Finally, variants within TTN or Mucin genes or variants exhibiting a gnomAD[10] minor allele frequency > 0.10 were removed.

Mutational significance was determined using MuSiC[11] with default settings on the filtered variant list. Potential cancer genes were prioritized via MUFFINN[12] using the HumanNet V1 network and the ndmax method. Genes with a probabilistic score > 0.5 were selected as candidate cancer genes of interest.

**DNA Mutational Signatures**

The R library deconstructSigs[13] was used to identify patterns in the somatic variants and their surrounding nucleotide context that support specific DNA mutational signatures in each sample. The somatic variants of each sample were compared to version 2 of the COSMIC mutational signature reference, which contains 30 unique signatures.[14] Briefly, a final variant list was constructed as described above except the gnomAD allele frequency filter was not applied and synonymous variants were included in the final variant list for this analysis. Samples with less than 45 total nonsynonymous SNVs (N=8) were omitted from the analysis. The “exome2genome” normalization method within deconstructSigs was used to obtain COSMIC signature weights for each sample.

**Copy Number Variant Calling**

Tumor ploidy aberrations were identified via cnvkit (0.9.6),[15] from the somatic_exome pipeline described above. A somatic amplification or deletion was defined from the segmented log2 tumor/normal ratio output from cnvkit as +/- 0.5. Copy number plots were made with the R library GenVisR (1.18.1)[16]. Significant amplified or deleted regions were identified with GISTIC (2.0.23)[17] using default parameters.

**References**

1. Griffith, M. *et al.* Genome Modeling System: A Knowledge Management Platform for Genomics. *PLoS computational biology* **11**, e1004274 (2015).

2. Auwera, G. A. V. der *et al.* From FastQ Data to High-Confidence Variant Calls: The Genome Analysis Toolkit Best Practices Pipeline. *Current Protocols in Bioinformatics* **43**, 11.10.1-11.10.33 (2013).
3. Saunders, C. T. *et al.* Strelka: accurate somatic small-variant calling from sequenced tumor-normal sample pairs. *Bioinformatics* **28**, 1811–1817 (2012).
4. Cibulskis, K. *et al.* Sensitive detection of somatic point mutations in impure and heterogeneous cancer samples. *Nature Biotechnology* **31**, 213–219 (2013).
5. Koboldt, D. C., Larson, D. E. & Wilson, R. K. Using VarScan 2 for Germline Variant Calling and Somatic Mutation Detection. *Curr Protoc Bioinformatics* **44**, 15.4.1-17 (2013).
6. Ye, K., Schulz, M. H., Long, Q., Apweiler, R. & Ning, Z. Pindel: a pattern growth approach to detect break points of large deletions and medium sized insertions from paired-end short reads. *Bioinformatics* **25**, 2865–2871 (2009).
7. McLaren, W. *et al.* The Ensembl Variant Effect Predictor. *Genome Biology* **17**, 122 (2016).
8. Ainscough, B. J. *et al.* A deep learning approach to automate refinement of somatic variant calling from cancer sequencing data. *Nat. Genet.* **50**, 1735–1743 (2018).
9. Barnell, E. K. *et al.* Standard operating procedure for somatic variant refinement of sequencing data with paired tumor and normal samples. *Genet Med* **21**, 972–981 (2019).
10. Karczewski, K. J. *et al.* The mutational constraint spectrum quantified from variation in 141,456 humans. *Nature* **581**, 434–443 (2020).
11. Dees, N. D. *et al.* MuSiC: Identifying mutational significance in cancer genomes. *Genome Res* **22**, 1589–1598 (2012).
12. Cho, A. *et al.* MUFFINN: cancer gene discovery via network analysis of somatic mutation data. *Genome Biology* **17**, 129 (2016).
13. Rosenthal, R., McGranahan, N., Herrero, J., Taylor, B. S. & Swanton, C. deconstructSigs: delineating mutational processes in single tumors distinguishes DNA repair deficiencies and patterns of carcinoma evolution. *Genome Biology* **17**, 31 (2016).

14. Alexandrov, L. B. *et al.* Signatures of mutational processes in human cancer. *Nature* **500**, 415–421 (2013).

15. Talevich, E., Shain, A. H., Botton, T. & Bastian, B. C. CNVkit: Genome-Wide Copy Number Detection and Visualization from Targeted DNA Sequencing. *PLOS Computational Biology* **12**, e1004873 (2016).

16. Skidmore, Z. L. *et al.* GenVisR: Genomic Visualizations in R. *Bioinformatics* **32**, 3012–3014 (2016).

17. Mermel, C. H. *et al.* GISTIC2.0 facilitates sensitive and confident localization of the targets of focal somatic copy-number alteration in human cancers. *Genome Biology* **12**, R41 (2011).
